# COATS Identifies Copy-Number-Dependent Drivers and Enablers of Aneuploidy in Cancer

**DOI:** 10.64898/2026.06.22.733880

**Authors:** Gaojianyong Wang, Khanh N. Dinh, Fabio Alfieri, Somayeh Fani, Teresa Davoli

## Abstract

Aneuploidy is a hallmark of most cancers but at the same time has been shown to decrease cellular fitness. Thus, to solve this conundrum, a current hypothesis in the field is that specific SCNAs may promote tolerance to the aneuploid state and/or promote additional chromosomal instability (CIN) and more aneuploidy. In other words, gains or losses of oncogenes (OGs) and tumor suppressor genes (TSGs) can, in turn, drive further CIN, promoting additional somatic alterations, or enhance aneuploid cell survival. Despite their importance, CN-dependent OGs and TSGs associated with aneuploidy remain largely unidentified. Here, we present a new method, Copy-number-dependent Oncogenes And Tumor Suppressors (COATS), to identify pan-cancer and cancer-specific CN-dependent OGs and TSGs associated with aneuploidy (Aneu-OGs and Aneu-TSGs). COATS integrates information theory and statistical tests to analyze gene expression, copy number, and aneuploidy, and incorporates timing analysis to distinguish early drivers of CIN from late tolerance enablers. Interestingly, using the CINner simulation framework, we show that aneuploidy drivers tend to occur earlier than aneuploidy enablers. Applying COATS to 33 TCGA cancer types, we identified 479 pan-cancer amplification-dependent Aneu-OGs and 141 deletion-dependent Aneu-TSGs. For validation, we used shRNA to knock down CCT5, a COATS-identified pan-cancer Aneu-OG predicted to promote aneuploidy tolerance, in isogenic aneuploid and near-diploid cells. Strikingly, CCT5 depletion was selectively toxic in aneuploid cells, supporting its classification as an aneuploidy tolerance enabler gene. Overall, our study defines a set of CN-dependent genes associated with aneuploidy and points to candidate therapeutic targets for chromosomally unstable cancers.

## Introduction

Aneuploidy, defined as an abnormal number of chromosomes in a cell, is a hallmark of cancer and a major contributor to tumorigenesis [1–3]. It is closely linked to genomic instability and frequently results in somatic copy number alterations (SCNAs) involving entire chromosomes, chromosome arms, or focal DNA segments [4]. These SCNAs disrupt normal gene dosage and alter RNA and protein levels of oncogenes (OGs) and tumor suppressor genes (TSGs) within affected regions [5]. Aneuploidy levels rise progressively during cancer development, from low levels in normal tissues to higher levels in precancerous lesions, and the highest levels in advanced and metastatic tumors [3, 6, 7]. This accumulation reflects both ongoing selective pressures during tumor evolution and the cellular adaptations that enable survival in the aneuploid state. A fundamental question, however, remains unresolved: which specific SCNAs drive this chromosomal instability (CIN), and which ones confer tolerance to the aneuploid state itself?

Aneuploidy and CIN are reciprocally linked: CIN generates aneuploidy through chromosome missegregation, and the resulting aneuploid state can, in turn, exacerbate CIN, creating a feedback loop that drives tumor evolution [4, 8]. The roles of specific SCNAs in this cycle, however, remain largely unclear. Amplifications or deletions of genes involved in cell cycle (CC) control and mitosis can promote the accumulation of further genomic alterations, while other SCNAs that arise during tumor evolution are selectively maintained because they alter the dosage of genes conferring a fitness advantage in chromosomally unstable cells [9]. Critically, aneuploidy itself induces multiple cellular stresses reducing fitness under normal conditions, including proteotoxic stress, metabolic imbalance, and replication stress. This paradox suggests that certain cancer-associated SCNAs serve a dual function: they drive additional chromosomal alterations by promoting CIN while also enabling cells to tolerate the aneuploid state by alleviating these stresses (**Fig. 1A**). Distinguishing CIN drivers from tolerance enablers, and identifying SCNAs that serve both roles, is therefore essential for understanding how aneuploidy accumulates during tumorigenesis.

**Fig. 1.**
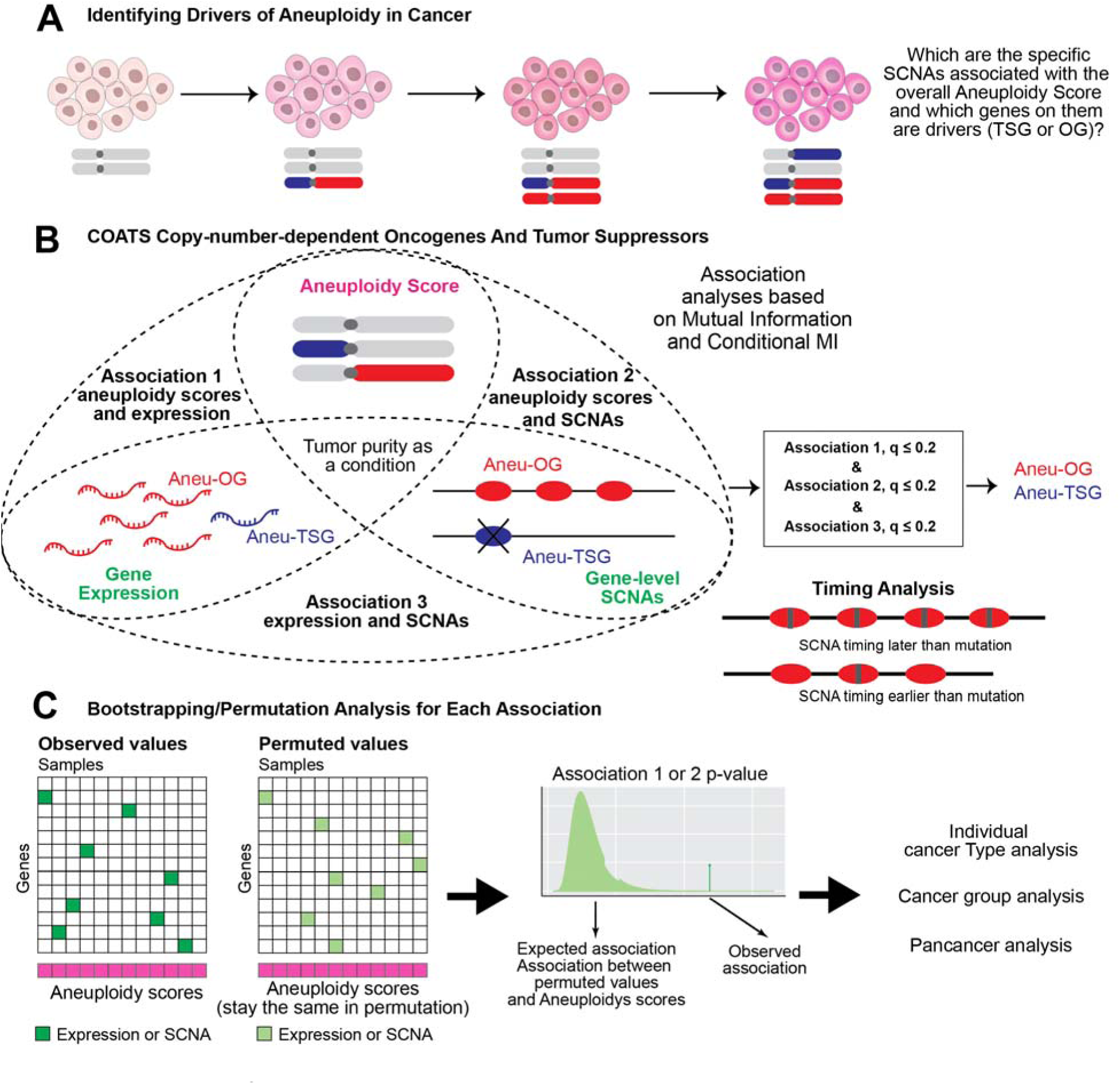
Overview of the COATS pipeline. **(A)** Schematic of aneuploidy progression in cancer cells. Cells begin with a normal karyotype and acquire SCNAs in CIN drivers and aneuploidy drivers, leading to chromosome missegregation and further SCNAs (gains in red, losses in blue). Accumulated SCNAs provide tolerance and fitness in highly aneuploid states. The aim of this study is to identify, among the SCNAs associated with aneuploidy, the OGs and TSGs that act as CIN drivers or as tolerance enablers. **(B)** Schematic of the COATS method for identifying CN-dependent Aneu-OGs and Aneu-TSGs. COATS evaluates three associations: Association 1, between AS and gene expression, conditioned on tumor purity; Association 2, between AS and SCNA levels, conditioned on tumor purity; and Association 3, between gene expression and SCNA levels. Mutual Information and Conditional Mutual Information are used to quantify these associations (Materials and Methods). Genes are classified as Aneu-OGs or Aneu-TSGs when all three associations are statistically significant (q ≤ 0.2) and show consistent directionality. Timing analysis then determines whether SCNAs occurred before or after somatic mutations, providing evidence as to whether each Aneu-OG or Aneu-TSG functions as a CIN driver or a tolerance enabler. **(C)** Bootstrapping and permutation approach used to evaluate the associations between gene expression or SCNAs and AS. “Observed values” show the actual expression or SCNA matrix; “Permuted values” show a shuffled expression or SCNA matrix in which AS remain constant. Permutation generates the null distribution against which observed associations are compared, yielding a p-value, and the false discovery rate (q-value) is computed from these p-values. The procedure is applied to individual cancer type, cancer group, and pan-cancer analyses.

In contrast to mutation-dependent OGs and TSGs associated with aneuploidy, which have been more extensively studied, CN-dependent OGs and TSGs associated with aneuploidy remain largely unidentified. The most prominent example among mutation-based drivers is TP53, whose loss of function permits the survival and proliferation of cells that would otherwise arrest or undergo apoptosis after chromosome missegregation [10]. By comparison, no systematic method exists to identify CN-dependent OGs and TSGs associated with aneuploidy or other cancer hallmarks. A recent study identified the top 5 genes whose mutations and the top 100 genes whose overexpression are associated with CIN [11], but expression-based analyses alone cannot distinguish whether overexpression reflects driver events that promote aneuploidy, consequences of pre-existing aneuploidy, or tolerance mechanisms enabling survival in the aneuploid state.

Existing approaches for separating tumor-driving SCNAs from passenger SCNAs fall into two categories: statistical tests for recurrent SCNAs, exemplified by GISTIC [12], and frequency-based analyses to identify significant alterations [13]. Because SCNA data alone are not sufficient to define functional OGs and TSGs, these methods are often complemented by integrated analysis of gene expression and copy number to refine candidate driver lists [5, 13]. Genome-wide association studies (GWAS) linking gene expression to cancer hallmarks have similarly been used to identify hallmark-associated genes, though such gene sets are not necessarily OGs or TSGs. The CIN70 signature, derived from this kind of analysis, is one such example [14].

Here, we present Copy-number-dependent Oncogenes And Tumor Suppressors (COATS), a method that integrates gene expression, SCNA, cancer hallmark, and mutation data to identify pan-cancer and cancer-specific CN-dependent OGs and TSGs and to infer their relative timing during tumor evolution. We apply COATS to 33 TCGA cancer types and their aneuploidy scores to identify amplification-dependent OGs associated with aneuploidy (Aneu-OGs) and deletion-dependent TSGs associated with aneuploidy (Aneu-TSGs) that either drive further CIN or enable tolerance to the aneuploid state. To validate the method experimentally, we used shRNA to deplete CCT5, a COATS-identified pan-cancer Aneu-OG predicted to enable aneuploidy tolerance, in isogenic near-diploid and aneuploid cells. CCT5 depletion was selectively toxic in aneuploid cells, supporting its role as an aneuploidy tolerance gene. Beyond aneuploidy, we demonstrate that COATS can be applied to identify CN-dependent OGs and TSGs associated with other cancer hallmarks, including immune and CC signatures.

## Results

### Overview of COATS, a new algorithm to identify SCNA-dependent oncogenes and tumor suppressors

We developed COATS (Copy-number-dependent Oncogenes And Tumor Suppressors), a computational method to identify genes whose SCNAs (gains or losses), together with the associated expression changes, are linked to specific cancer hallmarks or tumor features. Although the approach can be applied to any quantifiable tumor characteristic, we focused initially on aneuploidy, defined here as the total number of arm-level gains and losses, i.e., the aneuploidy scores (AS) [3]. We refer to the SCNA-dependent oncogenes and tumor suppressor genes associated with aneuploidy as Aneu-OGs and Aneu-TSGs, respectively.

COATS is based on the reasoning that for a true Aneu-OG or Aneu-TSG, the SCNA level, the expression level, and the aneuploidy score should all be jointly associated (**Fig. 1A**). For an Aneu-OG, gene gain should correspond to increased expression, and both the gain and the increased expression should be associated with higher AS. Conversely, for an Aneu-TSG, gene loss should correspond to decreased expression, and both the loss and the decreased expression should be associated with higher aneuploidy. COATS therefore evaluates three associations (**Fig. 1B**): (1) aneuploidy score versus gene expression, (2) aneuploidy score versus SCNA level, and (3) gene expression versus SCNA level. Because tumor purity [15] can confound the first two relationships, we include it as a confounding factor [16] (**Materials and Methods**). Statistical significance is assessed through a permutation-based method [16] (**Fig. 1C, Materials and Methods**).

A gene is classified as an SCNA-dependent OG or TSG only when all three associations are statistically significant (FDR q < 0.2 for each), and is further classified as an amplification-dependent Aneu-OG or a deletion-dependent Aneu-TSG only when the three associations show consistent directionality: all positive for Aneu-OGs and all negative for Aneu-TSGs (**Fig. 1B, Materials and Methods**). This dual requirement of statistical significance and concordant directionality is central to COATS, because it ensures that candidate Aneu-OGs and Aneu-TSGs reflect coordinated changes across SCNAs, expression, and aneuploidy rather than incidental correlations.

We applied COATS to publicly available TCGA data [17] covering 33 cancer types (**Materials and Methods**). COATS first identified cancer-specific SCNA-dependent Aneu-OGs and Aneu-TSGs (**Fig. 1C, Table S1**). To identify multi-cancer and pan-cancer associations, we then merged the results from individual cancer types using a meta-analytic approach (**Fig. 1C, Table S1, Materials and Methods**). We examined six cancer groups defined by tissue of origin or histological similarity: Adenocarcinoma Gastrointestinal (AdenoCarGI): COAD, READ, ESCA, STAD, PAAD; Adenocarcinoma Others (AdenoCarOther): BRCA, PRAD, LUAD, LIHC; Neural Crest Cells (Neural Crest): LGG, GBM, SKCM, UVM, THCA; Gynecological (CESC, UCEC, OV), Kidney (KICH, KIRC, KIRP), and Squamous (LUSC, HNSC, ESCA, BLCA, CESC). The pan-cancer analysis incorporated all 33 cancer types.

In summary, COATS identifies SCNA-dependent Aneu-OGs and Aneu-TSGs by integrating gene expression, SCNA, and aneuploidy data through three jointly evaluated associations, and operates at three resolutions: the individual cancer type, cancer group, and pan-cancer.

### Pan-cancer analysis reveals 479 Aneu-OGs and 141 Aneu-TSGs

Applying COATS to all 33 cancer types in TCGA (**Materials and Methods**), we identified 479 amplification-dependent pan-cancer Aneu-OGs and 141 deletion-dependent pan-cancer Aneu-TSGs (**Fig. 2A, Table S1**). Before applying the concordance requirement across the three criteria, a much larger number of genes satisfied the individual association criteria: expression of 871 and 653 genes respectively showed positive and negative associations with AS; SCNAs of 25,441 and 27,587 genes respectively showed positive and negative associations with AS; and 9,946 genes showed significant expression-SCNA associations. The requirement for concordance across all three criteria substantially narrows the candidate set and filters out genes whose expression changes are likely consequences of aneuploidy rather than drivers or tolerance enablers. The marked asymmetry between Aneu-OGs and Aneu-TSGs (479 vs. 141) is addressed in the Discussion.

**Fig. 2.**
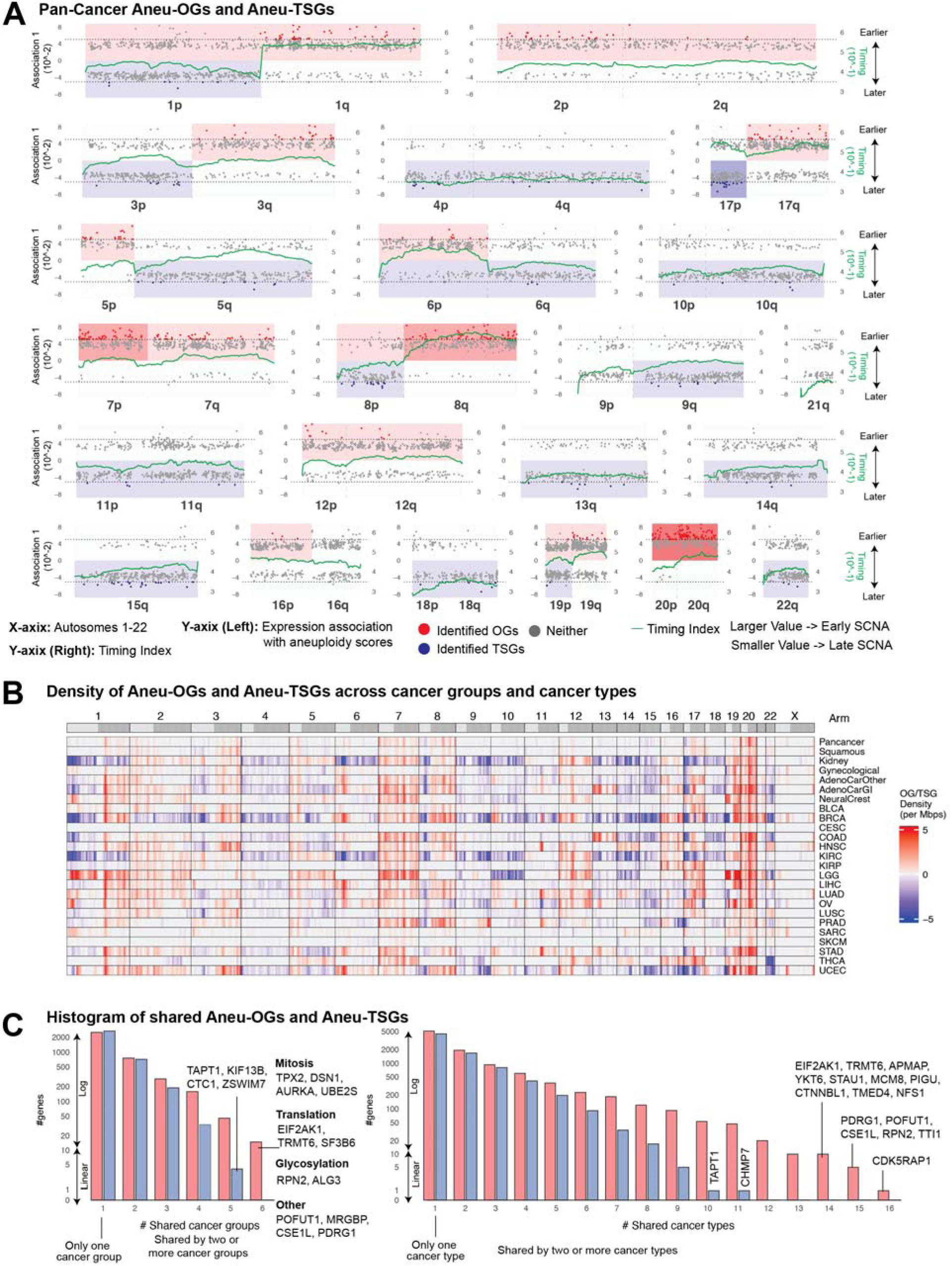
COATS identifies pan-cancer and tissue-specific Aneu-OGs and Aneu-TSGs. **(A)** Chromosomal mapping of pan-cancer Aneu-OGs and Aneu-TSGs. Each subfigure corresponds to one chromosome and shows the distribution and density of Aneu-OGs (red dots) and Aneu-TSGs (blue dots). Background shading represents Aneu-OG and Aneu-TSG density along the chromosome arm. The left y-axis shows the expression association with AS (Association 1 in Fig. 1), and the right y-axis (green line) shows the timing index (Materials and Methods). Higher timing index values denote earlier amplification events; lower values denote later events. **(B)** Genome-wide distribution of Aneu-OGs and Aneu-TSGs across cancer groups and cancer types. Each row represents a cancer group or cancer type. Aneu-OG and Aneu-TSG densities were calculated in 1 Mb genomic windows and plotted across chromosome arms (1–22 and X). Red indicates Aneu-OG density and blue indicates Aneu-TSG density; white indicates no Aneu-OGs or Aneu-TSGs. **(C)** Histograms of the number of Aneu-OGs (red) and Aneu-TSGs (blue) shared across cancer groups (left) and across cancer types (right). The x-axis indicates the number of cancer groups (left) or cancer types (right) in which a gene was identified as an Aneu-OG or Aneu-TSG.

The identified Aneu-OGs and Aneu-TSGs showed significant overlap with previously reported cancer gene sets while also including many previously unreported candidates. Among the 479 Aneu-OGs, 15 of them overlapped with the 226 amplification-dependent OGs from [5] (P < 1.2×10□, Fisher’s exact test), 6 of them overlapped with the 231 mutation-based OGs from [18] (P = 0.014, Fisher’s exact test), and 35 of them overlapped with the 100 aneuploidy-associated oncogenes from [11] (P < 2.2×10□^1^, Fisher’s exact test). Notably, the top 50 genes whose expression was positively associated with AS included 18 genes in the CIN70 signature [14] (P < 10□³, Fisher’s exact test). Among the 141 Aneu-TSGs, 9 genes overlapped with the 202 deletion-dependent TSGs from [5] (P < 2.4×10⁻□, Fisher’s exact test) and 5 genes overlapped with the 328 mutation-based TSGs from [18] (P = 0.0014, Fisher’s exact test). The limited intersection with mutation-based driver lists indicates that COATS identifies a largely distinct set of SCNA-dependent cancer genes.

Several well-established cancer genes appear among the Aneu-OGs and Aneu-TSGs (**Table S1**), providing further support for the method. CCNE1 (chr19), whose amplification is associated with poor prognosis across multiple cancer types, and E2F3 (6p), a transcription factor frequently overexpressed in cancer, were both identified as Aneu-OGs. TP53 (17p), the central regulator of genome stability, was identified as an Aneu-TSG, as was SMAD4 (18q), a core component of the TGF-β signaling pathway. We also observed genomic clustering of Aneu-OGs, including TPX2, E2F1, AURKA, and GNAS on chr20q, consistent with the frequent amplification of this region across cancer types.

In summary, the pan-cancer COATS analysis identified 479 Aneu-OGs and 141 Aneu-TSGs (**Fig. 2A, Table S1**) that overlapped significantly with established cancer gene sets while substantially expanding the catalog of SCNA-dependent genes associated with aneuploidy.

### Aneu-OGs are broadly shared across cancer types while Aneu-TSGs show tissue specificity

In addition to the pan-cancer analysis, we used COATS to identify Aneu-OGs and Aneu-TSGs for each individual cancer type and for the six cancer groups defined above: AdenoCarGI, AdenoCarOther, Neural Crest, Gynecological, Kidney, and Squamous (**Fig. 2B, Fig. S1, Table S1**). Across the six cancer groups, 220 Aneu-OGs were shared by at least four groups (more than half), while only 38 Aneu-TSGs met the same criterion. With a stricter threshold of at least five groups, 61 Aneu-OGs remained shared, compared with only 4 Aneu-TSGs (**Fig. 2C**). This asymmetry suggests that Aneu-OGs reflect common mechanisms of CIN and aneuploidy tolerance across cancer types, whereas Aneu-TSGs are more tissue specific, potentially reflecting distinct tumor suppression pathways or adaptive tolerance mechanisms in different cellular contexts (**Fig. S2A, Fig. S2B**).

A similar pattern emerged at the level of individual cancer types. Of the 33 cancer types analyzed, 23 had sufficient sample sizes for COATS to identify statistically significant Aneu-OGs and Aneu-TSGs (**Table S1**). Among these, 46 Aneu-OGs were shared by at least 12 of the 23 cancer types (more than half), whereas no Aneu-TSGs reached this threshold (**Fig. 2C**). The conservation of specific Aneu-OGs across diverse cancer types suggests that these genes represent core requirements for aneuploid and may serve as candidate targets for broad-spectrum therapeutic strategies. Altogether, Aneu-OGs converge on a shared set of genes broadly required across cancer types, whereas Aneu-TSGs are largely tissue specific, defining two distinct modes through which SCNAs contribute to aneuploidy in cancer.

### Enrichment of Aneu-OGs in cell cycle and stress response pathways

We performed GSEA using the identified Aneu-OGs and Aneu-TSGs to characterize their functional roles. Pan-cancer Aneu-OGs were strongly enriched in genes related to CC (**Fig. 3A**), consistent with the intrinsic link between CC dysregulation and the mitotic errors that generate aneuploidy, and with recent findings by Watson et al. [19]. This enrichment may reflect either a requirement for upregulated CC genes to sustain proliferation in cells with high CIN, or selection for SCNAs in these genes during tumor evolution. Beyond CC genes, pan-cancer Aneu-OGs were significantly enriched in mitosis, RNA metabolism, DNA replication, and DNA repair pathways (**Fig. 3A, Table S2**). We also identified several pan-cancer Aneu-OGs involved in epigenomic regulation, including DNMT3A, DNMT3B, EZH2 [20], CBX3, ACTL6A, ZNF273, POLR2K, SUV39H1, ZNF93, and TAF1A, suggesting that epigenetic alterations may also contribute to aneuploidy tolerance. In contrast, Aneu-TSGs did not show statistically significant enrichment in any pathway.

**Fig. 3.**
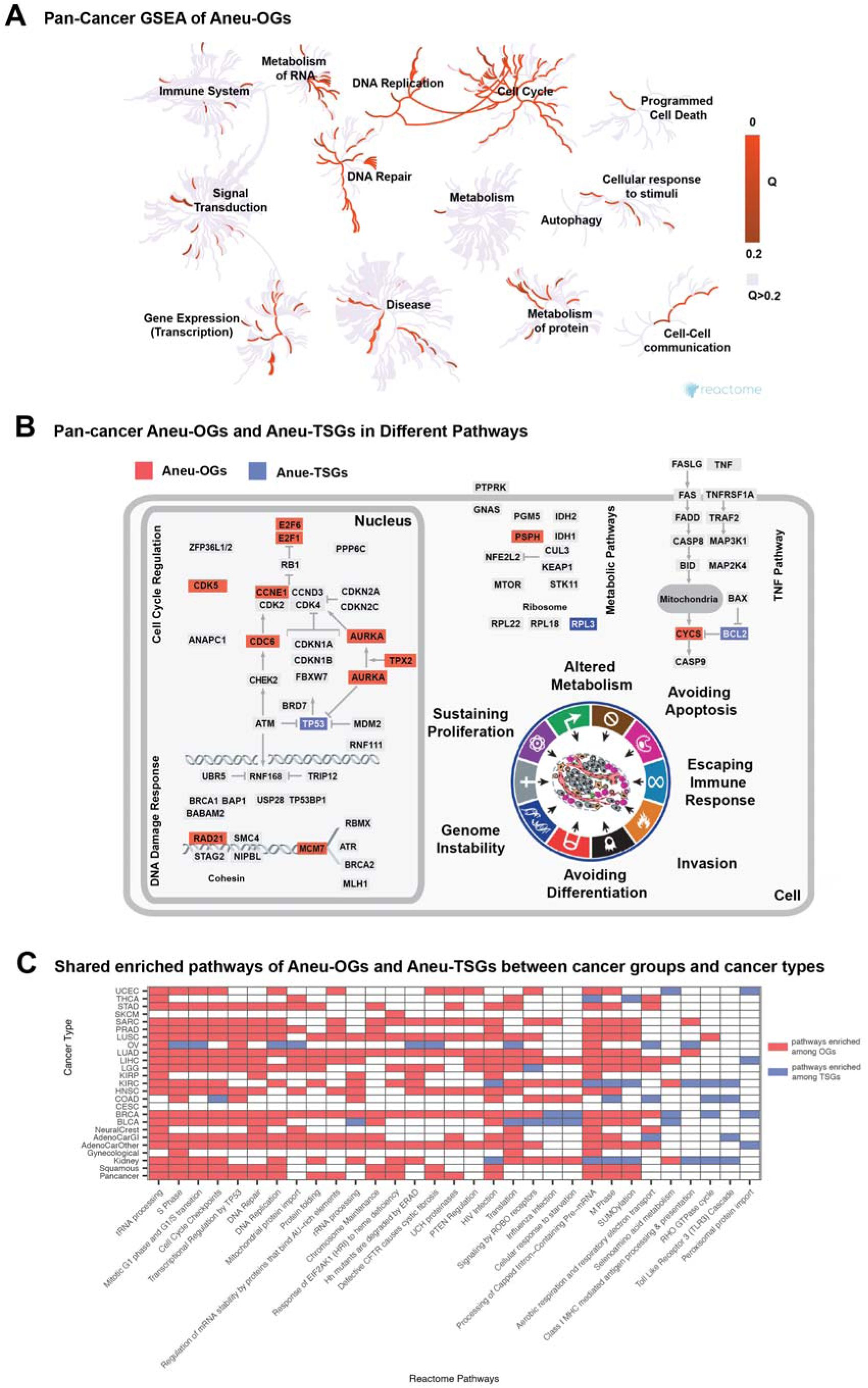
Pathways enriched among Aneu-OGs and Aneu-TSGs. **(A)** GSEA network of pan-cancer Aneu-OGs (Reactome; https://reactome.org). Each node represents a Reactome pathway, and connections indicate parent-subpathway relationships. Red color intensity reflects enrichment significance (Q value, 0 to 0.2), with darker red indicating stronger enrichment; pathways with Q > 0.2 are shown in gray. Pan-cancer Aneu-OGs were broadly enriched in CC, DNA replication, DNA repair, and RNA metabolism pathways, among others. **(B)** Pan-cancer Aneu-OGs (red) and Aneu-TSGs (blue) mapped onto cancer pathways associated with the hallmarks of cancer. Pathways shown include CC regulation and DNA damage response (associated with sustaining proliferation and genome instability), metabolic pathways (altered metabolism), and the TNF pathway (avoiding apoptosis). Selected representative Aneu-OGs and Aneu-TSGs from each pathway are labeled. **(C)** Heatmap of enriched pathways shared across cancer types and cancer groups. Each row represents a cancer type or group, and each column represents a Reactome pathway. Red cells indicate pathways enriched among Aneu-OGs, blue cells indicate pathways enriched among Aneu-TSGs, and white cells indicate pathways with no significant enrichment.

Pan-cancer Aneu-OGs were also linked to pathways involved in the DNA damage response, including RAD21 and MCM7, metabolism, including PSPH and RPL3, and TNF signaling, including BCL2 and CYCS (**Fig. 3B**). These findings indicate that aneuploidy is not achieved through alterations in a single pathway but rather through coordinated changes across multiple interconnected processes that govern cell proliferation, survival, DNA repair, apoptosis, and metabolic control.

Aneu-OGs from individual cancer types and cancer groups were enriched in the same pathways as pan-cancer Aneu-OGs, including CC, mitosis, DNA repair, and mitochondrial translation (**Fig. 3C**). The consistent enrichment in CC and mitosis pathways across cancer types reflects the universal role of disrupted cell division in cancer progression. Enrichment in DNA repair pathways suggests that aneuploid cancer cells require enhanced repair capacity to manage the genomic damage associated with chromosomal imbalance. Enrichment in mitochondrial translation pathways suggests that altered mitochondrial function, including energy production and apoptosis regulation, may support the metabolic demands of aneuploid cancer cells.

Altogether, Aneu-OGs converge on a coordinated set of pathways spanning CC progression, mitosis, DNA replication and repair, RNA metabolism, and mitochondrial function, indicating that aneuploidy is supported by the amplification of multiple interconnected biological programs rather than a single dominant mechanism.

### Simulations establish that the timing of SCNAs distinguishes aneuploidy drivers from enablers

Beyond identifying Aneu-OGs and Aneu-TSGs, we sought to infer the relative timing of these alterations during tumor evolution, as this would provide insight into their functional roles. We reasoned that SCNAs arising early in tumor development may actively promote CIN and drive the accumulation of aneuploidy (aneuploidy drivers), whereas SCNAs arising later primarily enable tolerance to the aneuploid state or confer growth advantages to cells that have already acquired aneuploid karyotypes (aneuploidy enablers).

To test whether the timing of SCNAs can, in principle, distinguish the functional roles of driver and enabler, we first performed an *in silico* study using a modified version of CINner [21], a simulator of CIN. We simulated copy number profiles under two scenarios. In the “CIN driver” setting, 5p gain increased the missegregation rate while CIN tolerance was held constant across cells. In the “CIN enabler” setting, 5p gain increased the CIN tolerance threshold for cell viability while the missegregation rate was held constant. Parameters were chosen so that the overall CIN scores (total missegregation counts) were comparable across different copy numbers of 5p in the two scenarios (**Fig. 4A**). Across 10,000 simulations of each scenario, we observed similar copy number profiles (**Fig. S3A, Fig. S3B**) and similar distributions of the real time at which 5p alterations occurred (**Fig. S3C, Fig. S3D**). However, the SCNA timing distribution, defined by the number of missegregations occurring before and after each 5p event, showed that 5p alterations arose significantly earlier in tumor evolution in the “CIN driver” scenario than in the “CIN enabler” scenario (**Fig. 4B**). These simulations confirm that the relative timing of an SCNA carries information about whether it acts as a driver or an enabler of aneuploidy, providing a conceptual basis for using timing to distinguish the two roles in real tumor data.

**Fig. 4.**
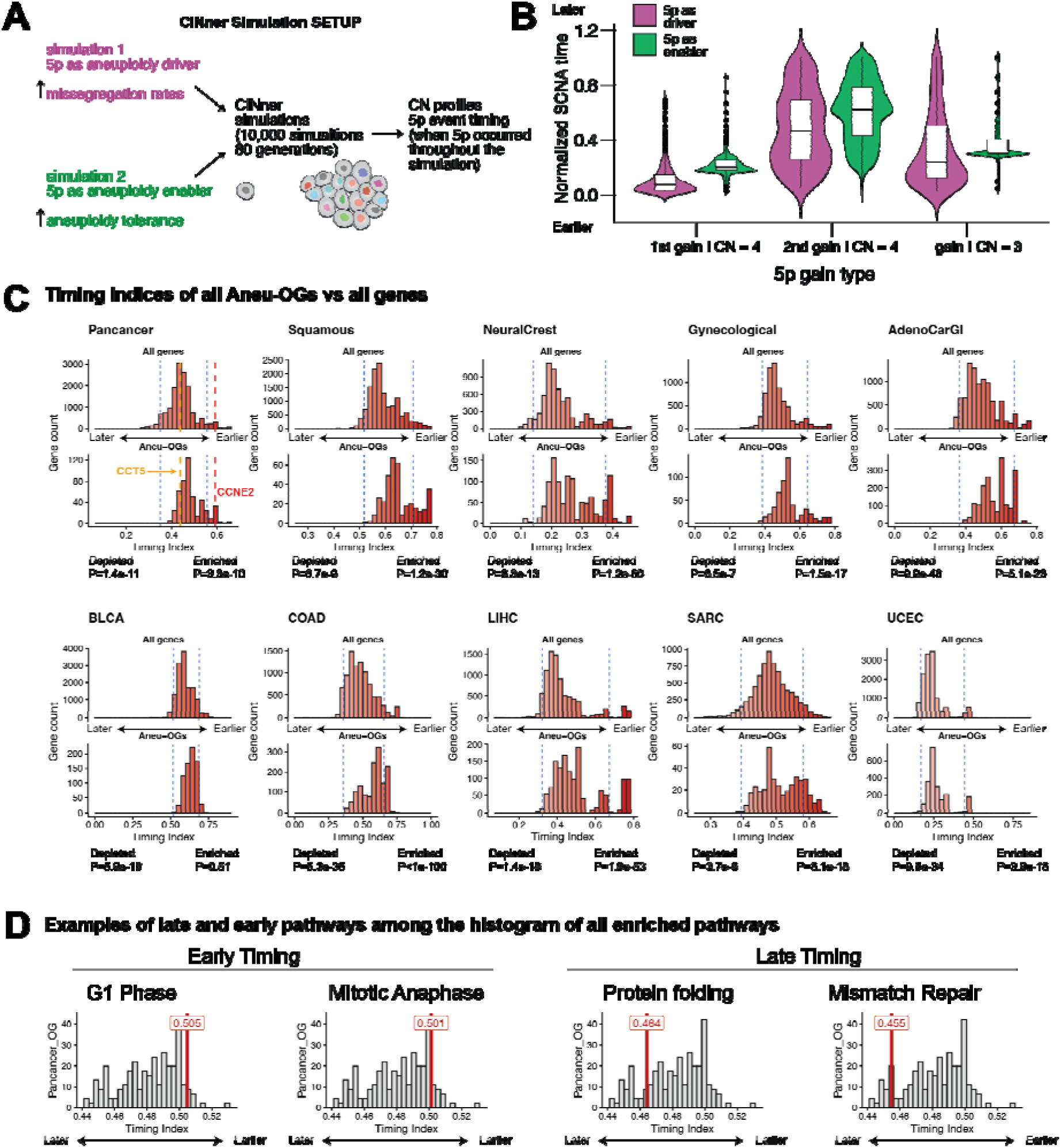
Timing analysis of Aneu-OGs and associated pathways. **(A)** Schematic of our simulation study comparing SCNA events as aneuploidy drivers versus aneuploidy enablers, illustrated using arm 5p. Parameters in each setting were calibrated to yield comparable CIN scores across synthetic samples. Simulated data are compared to identify observable differences between the two scenarios for the role of arm 5p. **(B)** Comparison of normalized SCNA timing for 5p SCNA events between the two scenarios shown in (A). SCNAs are grouped into three categories: exclusive 5p gains resulting in observed CN = 3, and first or second 5p gains resulting in observed CN = 4. 5p events consistently arise earlier in SCNA time across all categories when acting as aneuploidy drivers than as enablers. **(C)** Timing index distributions for all genes compared with Aneu-OGs across ten cancer groups or cancer types. In each panel, the upper histogram shows the unique timing indices of all genes and the lower histogram shows the timing indices of Aneu-OGs. The timing indices were classified as early, intermediate, or late events using the 5th and 95th percentile thresholds of the timing indices of all the genes, indicated by the blue dashed lines. Compared with the unimodal distribution observed for all genes, Aneu-OGs show more complex timing distributions in several cancer groups and types. Aneu-OGs tend to be enriched in early events but depleted in late events. The number of late-timing Aneu-OGs are more than that of the early-timing Aneu-OGs. The timing index for CCT5 (orange) and CCNE2 (red) are marked in Pan-cancer. **(D)** Timing index (red vertical line) of three representative Reactome pathways enriched among pan-cancer Aneu-OGs, plotted against the histogram of timing indices for all enriched pan-cancer Aneu-OG pathways: G1 Phase (R-HSA-69236, left), Mitotic Anaphase (R-HSA-68882, middle), and Protein folding (R-HSA-391251, right).

### Timing analysis in human tumors separates aneuploidy drivers from enablers

To estimate the timing of SCNAs in TCGA tumors, we leveraged somatic mutation data from the Multi-Center Mutation Calling in Multiple Cancers [22]. The underlying principle is that if a mutation occurs before an amplification event, the mutation will be duplicated along with the amplified region and thus be present in all copies [23, 24]. Conversely, if amplification precedes the mutation, only a subset of the amplified copies will harbor the mutation. By analyzing the distribution of mutations across amplified regions, we inferred the chronological order of SCNA events and assigned a timing index ranging from 0 to 1, where larger values indicate earlier events and smaller values indicate later events (**Fig. 1B, Materials and Methods**).

We observed chromosome arm level heterogeneity in the relative timing of amplifications. For instance, in our pan-cancer analysis (**Fig. 2A**), amplification of 8q occurred earlier than that of 8p. The timing index can also vary within a single chromosome arm: amplification of 5p and 5q were generally late, with the exception of 5q31–q32. Amplification of chr17 was generally an early event, while amplifications of 5p, chr4, and chr10 tended to occur later. Relative timing also varied across cancer groups. For example, in Kidney, amplifications of 3p and 5q were early events (**Fig. S1**), whereas in Neural Crest, these same amplifications occurred relatively late (**Fig. S1**). Chr13 amplification was early in AdenoCarGI but late in the other five cancer groups (**Fig. S1**). These findings indicate that different regions of the same chromosome, and the same chromosome arm across different tumor contexts, can be amplified at distinct stages of tumorigenesis. This temporal heterogeneity likely reflects the differential timing at which chromosome arms, and the molecular pathways they encode, contribute to survival advantages and promote CIN in different cancer types.

We next compared the distribution of timing indices of the Aneu-OG with the timing distribution of all genes (**Fig. 4C**), which appear unimodal and normally distributed (P=2.2×10^-16^, Shapiro–Wilk test in pan-cancer). Within each cancer type or cancer group, timing indices were classified into three empirical groups (**Fig. 4C**): early events (high timing indices ≥ the 95th percentile), intermediate events (timing indices between the 5th and 95th percentile), and late events (low timing indices ≤ the 5th percentile). Several observations were evident when comparing these distributions. First, compared with the timing distribution of all genes, overall Aneu-OGs tended to be enriched in early events and depleted in late events (**Fig. 4C, Table S3**). For example, among all the identified pan-cancer Aneu-OGs, 58 were early events (enrichment P value = 3.3×10^-10^, hypergeometric test), none was late events (depletion P value = 1.4×10^-11^, hypergeometric test), and 421 were intermediate events (enrichment P value = 0.95, depletion P value = 0.07, hypergeometric test). In other words, relative to the genome wide background, Aneu-OGs were shifted toward earlier timing, with a minority reaching the earliest timing and the majority occupying intermediate timing values (**Fig. 4C**).

Second, compared with the approximately unimodal timing distribution observed for all genes, Aneu-OGs displayed bimodal distributions in multiple cancer types (**Fig. 4C**), with a smaller population at early timing (high timing indices ≥ the 95th percentile) and a larger population at intermediate timing (timing indices between the 5th and 95th percentile). This distribution was most consistent with a small set of Aneu-OGs acting as drivers of aneuploidy that arose earlier than most of the genes in the whole-genome, alongside a larger set of relatively later occurring Aneu-OGs that likely acted as enablers of aneuploidy tolerance.

Altogether, relative to the unimodal genome-wide background, Aneu-OGs were shifted toward earlier timing, but their distribution was bimodal, comprising a smaller population of early-arising drivers of aneuploidy and a larger population of intermediate-timing Aneu-OGs that likely act as enablers of aneuploidy tolerance.

### Pathway level timing distinguishes pathways that drive versus enable aneuploidy

We next assigned a timing index to each enriched Aneu-OG pathway in each cancer type and cancer group (**Table S2, Materials and Methods**). The timing index of a pathway is the average of the timing indices of its enriched Aneu-OGs. Two cancer types or groups may show enrichment for the same pathway, but the specific Aneu-OGs contributing to that enrichment can differ between them. Timing analysis of enriched Aneu-OG pathways revealed recurrent early involvement of CC and mitosis pathways across cancer types and groups. In the pan-cancer analysis, pathways related to the G1/S transition, G1 phase, and mitotic anaphase showed relatively high timing indices (**Fig. 4D, Table S2**), indicating that these transcriptional and functional programs are associated with driving aneuploidy in cancer. A similar pattern was observed in individual cancer types and cancer groups: G1 phase (R-HSA-69236) was early in AdenoCarGI, BLCA, and SARC (**Fig. S2D, Table S2**), and mitotic anaphase (R-HSA-68882) was early in AdenoOther, BRCA, LIHC, SARC, and UCEC (**Fig. S2E, Table S2**). These results indicate that the generation of aneuploidy recurrently involves CC regulation across diverse cancer contexts.

In contrast, Aneu-OG enrichments in protein metabolism pathways occurred as late events (**Fig. 4D, Fig. S2F, Table S2**). The timing index for the Protein folding pathway (R-HSA-391251) was low in pan-cancer, Kidney, LGG, and LUSC (**Fig. S2F, Table S2**). Within Protein folding, the Folding of actin by CCT/TRiC subpathway was also late (**Fig. 5A**) and includes CCT5, which lies on the late amplified 5p (**Fig. 2A**, **Fig. 4C**). Altogether, pathway level timing analysis identified CC and mitosis programs, including G1/S transition, G1 phase, and mitotic anaphase, as the earliest and most recurrent driver associated pathways across cancer contexts, whereas protein folding pathways, including the CCT/TRiC chaperonin complex, arose late and were associated with tolerance to the aneuploid state.

**Fig. 5.**
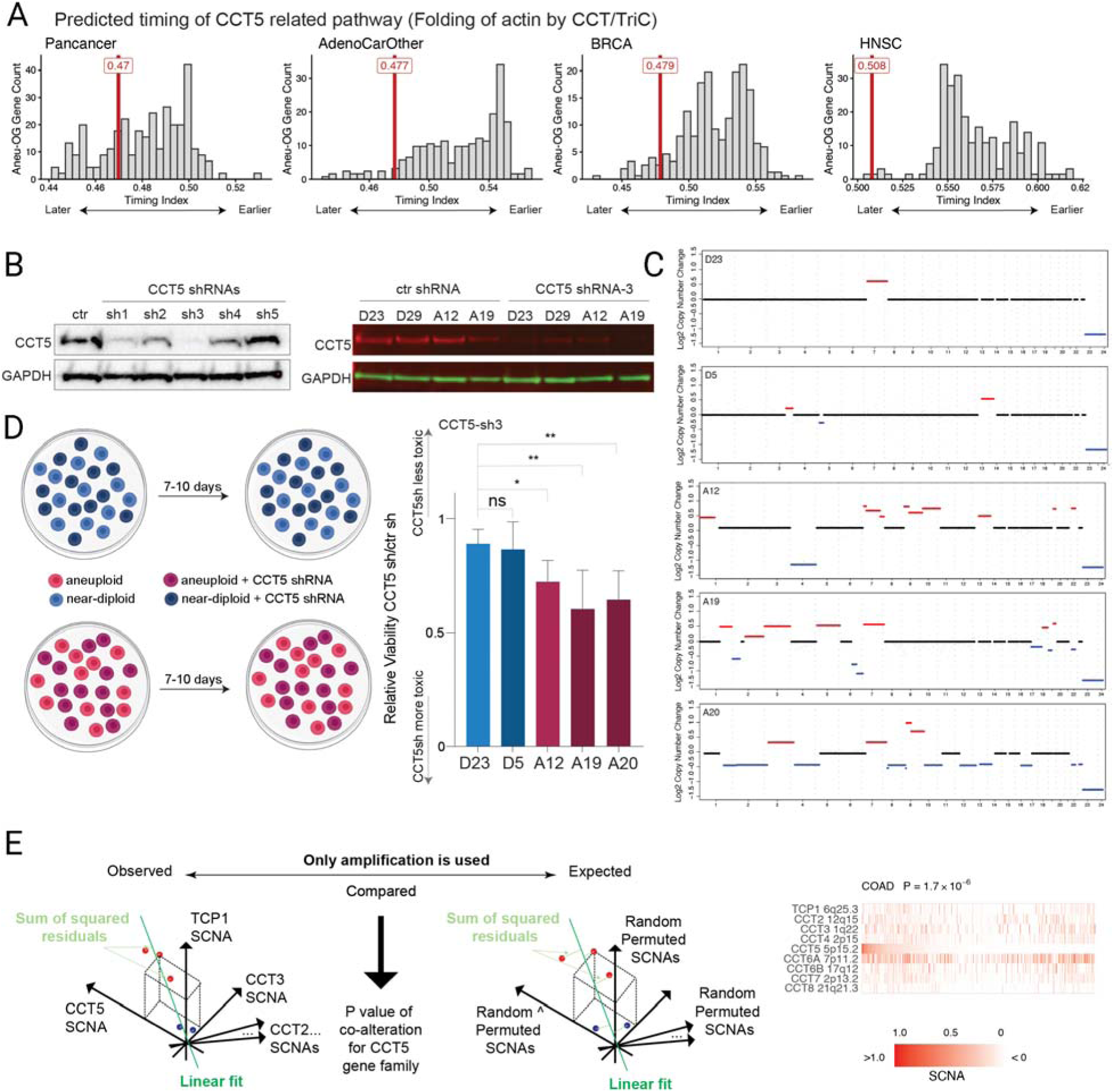
CCT5 is selectively essential in aneuploid HCEC clones. **(A)** Timing index (red vertical line) of the Reactome pathway “Folding of actin by CCT/TRiC” (R-HSA-390450) plotted against the histogram of timing indices for all enriched Aneu-OG pathways across cancer groups and cancer types. **(B)** Western blot analysis of CCT5 and GAPDH protein levels in cells transduced with non-targeting control shRNA or with shRNAs targeting human CCT5. The left panel shows the parental HCEC clone treated with five different CCT5-targeting shRNAs (sh1–sh5). The right panel shows the indicated HCEC clones (D23, D29, A12, A19) transduced with control shRNA or CCT5 shRNA-3. **(C)** Copy number profiles of the indicated HCEC clones obtained by low-pass whole-genome sequencing. Red segments indicate genomic gains and blue segments indicate genomic losses. Representative near-diploid (D23, D5) and aneuploid (A12, A19, A20) clones are shown. Chromosome numbers are indicated on the x-axis. **(D)** *(Left)* Schematic of the multicolor competition assay. Near-diploid or aneuploid HCEC clones transduced with control shRNA or CCT5 shRNA-3 were mixed at a 1:1 ratio and co-cultured for 7 to 10 days; the relative proportions of each population were then quantified by flow cytometry, with one population fluorescently labeled and the other unlabeled. Experiments were performed in triplicate with fluorescent labels switched between replicates. *(Right)* Quantification of the multicolor competition assay for the indicated near-diploid (D23, D5) and aneuploid (A12, A19, A20) HCEC clones, expressed as the relative viability of CCT5 shRNA-expressing cells compared with control shRNA-expressing cells. Statistical significance was assessed by Wilcoxon rank-sum test (ns, not significant; *P < 0.05; **P < 0.01). Error bars represent standard deviation from three independent experiments. **(E)** Analysis of SCNA co-amplification in the CCT gene family (TCP1, CCT2, CCT3, CCT4, CCT5, CCT6A, CCT6B, CCT7, and CCT8) across cancer types. The top schematic illustrates the method: observed amplifications of the CCT gene family are fit linearly in a multidimensional space (with all deletions set to zero) and the sum of squared residuals is calculated. The observed value is compared against a null distribution obtained by random permutation of the amplifications to derive a p-value for co-alteration of the CCT family. The heatmaps (bottom) show individual amplifications for the CCT family genes in COAD, KICH, LAML, LIHC, PCPG, PRAD, READ, and UVM. Color scale: red indicates amplification, with intensity representing the degree of amplification; SCNA values below zero are shown in white. Gene names and their cytogenetic locations are labeled on the left of each heatmap.

### CCT5 is an enabler of the aneuploid state

CCT5 was identified by COATS as a pan-cancer Aneu-OG with later timing (**Fig. 2A**, **Fig. 4C, Fig. S2C**), and the later timing of pathway Folding of actin by CCT/TRiC (**Fig. 5A**), together with the later timing of 5p amplification (**Fig. 2A, Fig. S1**), suggests that CCT5 functions as a tolerance enabler rather than as an early driver of CIN. CCT5 is a subunit of the chaperonin-containing TCP-1 (CCT) complex, an essential molecular chaperone required for the proper folding of numerous cellular proteins, including cytoskeletal components such as actin and tubulin. Because aneuploid cells experience elevated proteotoxic stress arising from imbalanced protein stoichiometry, we hypothesized that increased CCT5 expression driven by 5p gain supports the protein folding capacity required to tolerate the aneuploid state.

To test whether CCT5 is preferentially required for aneuploid cell survival, we used an isogenic experimental system derived from human colon epithelial cells (HCECs) with inactivated TP53 [25]. This system allows direct comparison of CCT5 depletion in near-diploid versus aneuploid contexts in a controlled genetic background. We classified clones with one or two chromosome arm-level alterations as near-diploid, and those with more than five arm-level alterations as aneuploid (**Fig. 5C**).

We first screened five shRNAs targeting CCT5 in the parental HCEC clone using Western blot analysis (**Fig. 5B, left panel**) and selected CCT5 shRNA-3, which produced robust and consistent knockdown, for all subsequent experiments. CCT5 shRNA-3 efficiently depleted CCT5 protein in both near-diploid (D23, D29) and aneuploid (A12, A19) clones (**Fig. 5B, right panel**).

To quantitatively measure the differential requirement for CCT5 in near-diploid versus aneuploid cells, we performed a multicolor competition assay (**Fig. 5D, left panel**). Cells transduced with control shRNA or CCT5 shRNA-3 were differentially labeled, with one population expressing GFP and the other left unlabeled, and mixed at an initial 1:1 ratio. Mixed populations were co-cultured for 7 to 10 days, allowing multiple rounds of cell division, after which the final ratio of the two populations was measured by flow cytometry. A decrease in the proportion of CCT5 shRNA cells relative to control shRNA cells indicates that CCT5 depletion impairs competitive fitness. To control for any effects of fluorescent protein expression on cell fitness, each experiment was performed in triplicate with the fluorescent labels swapped between conditions across replicates.

Near-diploid clones (D23, D5) showed no significant change in the relative abundance of CCT5 shRNA cells compared with control shRNA cells, indicating that CCT5 knockdown does not compromise their competitive fitness (**Fig. 5D, right panel**). In contrast, all three aneuploid clones tested (A12, A19, A20) showed a significant reduction in the relative viability of CCT5 shRNA cells relative to controls (Wilcoxon rank-sum test, P < 0.01), demonstrating that CCT5 depletion confers a substantial competitive disadvantage specifically in aneuploid cells (**Fig. 5D, right panel**). This context-specific essentiality is consistent with our computational prediction and supports the classification of CCT5 as an aneuploidy tolerance enabler gene. The preferential requirement for CCT5 in aneuploid cells likely reflects their heightened proteotoxic stress and increased dependence on protein quality control machinery.

To determine whether this functional dependence is also reflected at the genomic level, we asked whether copy number gains affecting members of the CCT gene family tend to co-occur across human cancers. We applied a linear fit approach (**Fig. 5E, Materials and Methods**) to evaluate the co-amplification of TCP1, CCT2, CCT3, CCT4, CCT5, CCT6A, CCT6B, CCT7, and CCT8. In brief, observed amplifications of the nine genes are fit linearly in a multidimensional space (deletions are set to zero) and the sum of squared residuals (SSR) is computed; smaller SSRs indicate stronger co-occurrence. A null distribution of SSRs is generated by random permutation of the amplifications across samples, and the p-value reflects how the observed SSR compares with the null distribution. Co-amplification of CCT family genes was significantly more frequent than expected by chance in eight cancer types: COAD, KICH, LAML, LIHC, PCPG, PRAD, READ, and UVM (**Fig. 5E, Fig. S3E**).

Altogether, CCT5 depletion is selectively toxic in aneuploid cells, validating the COATS prediction that CCT5 acts as an aneuploidy enabler, and the recurrent co-amplification of multiple CCT subunits across cancers suggests that the entire CCT complex is under positive selection as a tolerance mechanism, identifying it as a candidate therapeutic target in chromosomally unstable cancers.

### OGs and TSGs associated with other cancer hallmarks

To demonstrate that COATS can identify OGs and TSGs associated with cancer hallmarks beyond aneuploidy, we applied the method to two additional signatures: an immune signature and a CC signature. The immune signature was defined as the average expression of CD247, CD2, CD3E, GZMH, NKG7, PRF1, and GZMK. The CC signature was defined as the average expression of CENPE, CCNA2, CCNB2, MCM6, CCNF, BUB1, CDC20, CDC6, CDK1, and PLK1. Pan-cancer and cancer-specific OGs and TSGs associated with immune (**Table S4**) and CC (**Table S5**) signatures were identified using the same statistical criteria as in the aneuploidy analysis, with the convention that Immune-TSGs are positively associated and Immune-OGs are negatively associated with the immune signature, reflecting the role of Immune-OGs in immune evasion.

No pan-cancer Immune-OGs or Immune-TSGs were identified (**Fig. S4A**). Across the six cancer groups, only two Immune-TSGs, TNFAIP8 [26] and NR3C1 [27], were shared by more than half of the groups (**Fig. S4B**), and 90 Immune-OGs and Immune-TSGs were shared by more than two groups (**Fig. S4B**). This tissue specificity is consistent with the idea that different tumors evolve distinct immune evasion strategies adapted to their cellular and microenvironmental contexts. Unlike the broadly shared Aneu-OG pathways, which converge on cell proliferation, differentiation, and survival, immune evasion appears to involve cancer type-specific SCNAs that do not generalize across tumor types. GSEA results support this interpretation: although enrichment of Immune-OGs and Immune-TSGs was identified within individual cancer types or groups (**Table S6**), no enriched pathway was shared across more than four cancer groups or cancer types (**Fig. S4C**).

For the CC signature, we identified 2,133 pan-cancer CC-OGs and 274 pan-cancer CC-TSGs (**Fig. S5A**), of which 339 CC-OGs and 51 CC-TSGs overlapped with the Aneu-OGs and Aneu-TSGs identified above. This is consistent with the universal role of CC disruption in tumorigenesis, where SCNAs affecting CC genes can promote both uncontrolled proliferation and CIN. A total of 288 CC-OGs and 18 CC-TSGs were shared by more than half of the six cancer groups (**Table S5**), with the shared CC-OGs likely reflecting a common mechanism of CC dysregulation across cancer types (**Fig. S5B**). Similar to Aneu-TSGs and Immune-TSGs, the smaller number of broadly shared CC-TSGs indicates that CC-TSG inactivation is also tissue specific. GSEA further supports these patterns (**Fig. S5C**): CC-OGs were enriched in mitosis, DNA replication, apoptosis, and mRNA splicing pathways across cancer groups and types (**Fig. S5C, Table S7**), whereas no CC-TSG enrichment was shared by more than four cancer groups or cancer types (**Fig. S5C, Table S7**).

Altogether, COATS successfully identifies SCNA-dependent OGs and TSGs associated with both immune and CC signatures, demonstrating its applicability beyond aneuploidy and revealing distinct sharing patterns across hallmarks: immune-associated SCNAs are largely tissue specific, whereas CC associated SCNAs are broadly shared across cancer types and substantially overlap with aneuploidy-associated genes.

## Discussion

COATS integrates gene expression, copy number, and aneuploidy data to prioritize candidate cancer genes whose genomic and transcriptomic profiles are most consistent with functional roles as Aneu-OGs or Aneu-TSGs. The method operates as a hypothesis-generating framework rather than a proof of causality, and the resulting gene lists represent high-confidence candidates that require experimental validation rather than definitive annotations. Several broad patterns emerge from applying COATS across 33 TCGA cancer types that offer insights into the biology of aneuploidy and CIN in cancer.

### Asymmetry and tissue specificity of Aneu-OGs versus Aneu-TSGs

A notable feature of the pan-cancer results is the numerical asymmetry between Aneu-OGs and Aneu-TSGs: we identified 479 amplification-dependent Aneu-OGs compared with only 141 deletion-dependent Aneu-TSGs. Beyond their numbers, Aneu-OGs are also more broadly shared across cancer types. Of the six cancer groups analyzed, 220 Aneu-OGs were shared by at least four groups, compared with only 38 Aneu-TSGs meeting that threshold. A total of 46 Aneu-OGs were identified in at least 12 of the 23 individual cancer types, whereas no Aneu-TSG reached this level of conservation (**Fig. 2**). This asymmetry is unlikely to reflect a technical bias, since the method applies identical statistical criteria to gains and losses. The predominance of amplification-driven over deletion-driven events echoes earlier pan-cancer copy number analyses showing that oncogene amplifications are more frequent and more broadly distributed across tumor types than tumor suppressor deletions [28]. The pattern likely reflects a genuine biological difference in how amplification-driven and deletion-driven mechanisms contribute to aneuploidy tolerance across cancer contexts: copy number gains converge on a common core of biological programs broadly required in aneuploid cancers, whereas Aneu-TSG losses appear more tissue-specific, potentially reflecting distinct tumor suppression pathways or adaptive tolerance mechanisms in different cellular contexts, consistent with what has been observed broadly for both copy-number-dependent and mutation-based TSGs [29, 30].

The kidney cancer group provides a striking illustration of this tissue specificity. The chromosomal arm-level alteration landscape in kidney cancers is, in several respects, inverted relative to that of other cancer groups: 1q, predominantly gained and enriched for Aneu-OGs in most other groups, shows loss in kidney cancers; 6p, typically lost in other groups, is gained; and 17q, broadly gained elsewhere, is deleted in kidney cancers (Fig. S1). At the pathway level, the M Phase pathway, enriched among Aneu-OGs in virtually all other cancer types and groups, is instead enriched among Aneu-TSGs in the kidney group (Fig. 3C). This is consistent with the known biology of clear cell renal cell carcinoma, in which 3p loss is the dominant and earliest genomic event, simultaneously inactivating one allele each of VHL, PBRM1, BAP1, and SETD2 [31]; PBRM1 and SETD2 both have direct roles in mitotic fidelity, with PBRM1 promoting chromosome cohesion [32] and SETD2 methylating tubulin to support proper spindle function [33]. The kidney group thus serves as an informative internal control, demonstrating that COATS captures cancer type-specific SCNA dependencies rather than a generic pan-cancer signal.

### Aneu-OGs converge on coordinated programs of genome maintenance and stress response

The functional identity of these broadly shared Aneu-OGs supports this interpretation. GSEA revealed consistent enrichment of mitosis, S phase, and DNA replication pathways across nearly all cancer types and groups analyzed (**Fig. 3**), including established regulators of mitotic entry and progression, such as AURKA, TPX2, UBE2C, NEK2, and BIRC5. The AURKA/TPX2 axis is of particular interest: both proteins are among the most frequently co-overexpressed mitotic regulators across human cancers, and their co-amplification has been shown to induce chromosome missegregation and attenuate p53 signaling in non-transformed cells, linking their gain not only to tolerance of an existing aneuploid state but also to the active generation of further CIN [34, 35]. TPX2 displays the strongest correlation with CIN among genes in the CIN70 signature [36], and its appearance as a pan-cancer Aneu-OG here is consistent with a dual role as both a driver and tolerance enabler depending on its timing. Amplification of S phase regulators, including MCM2, MCM4, MCM7, MCM8, GINS1, and TOP2A, likely supports both the replication demands of rapidly proliferating aneuploid cells and tolerance to the elevated replication stress in cells carrying unbalanced genomes. Aneu-OGs were also enriched in DNA repair pathways, including homologous recombination and mismatch repair genes, such as RAD51AP1, EXO1, and MSH2, suggesting that aneuploid cancer cells face increased DNA damage burdens and must amplify their repair capacity in parallel to maintain viability.

Beyond these canonical genome maintenance functions, Aneu-OGs are substantially enriched in RNA metabolism pathways, including mRNA splicing (SNRPB, SNRPC, MAGOHB, SF3B4, LSM5), RNA modification (NSUN2, BUD23, TRMT6), and RNA polymerase III transcription. A mechanistic basis for this dependency has recently been established. Proper splicing of key mitotic and cohesion regulators is itself required for chromosomal stability, as defective splicing of sororin disrupts sister chromatid cohesion [37]. In addition, aneuploid cells face a stoichiometric transcriptional burden due to their extra chromosomes and must upregulate RNA degradation pathways, including nonsense-mediated decay and miRNA-mediated silencing, to compensate for excess transcriptional output, rendering them selectively sensitive to perturbation of RNA processing capacity [38]. Thus, amplification of RNA metabolism genes among our Aneu-OGs may reflect the same adaptive logic: by increasing their capacity for accurate RNA processing and turnover, aneuploid cells manage the transcriptional imbalance that accompanies changes in genomic dosage. Finally, enrichment in protein folding and chaperonin pathways, including multiple CCT complex subunits (CCT3, CCT5, CCT6A) and proteasome components (PSMD4, PSMD11, PSMA7), is consistent with the proteotoxic stress that aneuploid cells experience as a consequence of stoichiometric imbalances among protein complex subunits [39, 40]. Together, these enrichments describe aneuploid cancer cells as having amplified not a single survival mechanism, but rather a coordinated set of programs spanning mitosis, replication, DNA repair, RNA metabolism, and protein homeostasis.

### Aneuploidy drivers are outnumbered by enablers

Our analysis reveals that a small set of Aneu-OGs occur early in tumor evolution, earlier than genes across the genome and earlier than CC-OGs and Immune-OGs. The enrichment of these Aneu-OGs in early timing suggests that their gains are not simply byproducts of CIN but are actively selected to establish and expand aneuploid cells. If these gains were passive consequences of CIN, their timing would be expected to follow the genome-wide timing distribution rather than being preferentially observed early; the early enrichment therefore points to positive selection during early tumor evolution, contributing to the initiation or early expansion of aneuploid cancer cells. Our simulations provide an interpretive framework for this signal: events arising early tend to act as drivers of aneuploidy, whereas events arising later tend to act as enablers that confer tolerance to the aneuploid state. We note that early and late timing are relative designations rather than absolute thresholds, so the boundary between drivers and enablers is not sharply defined. Even allowing for this caveat, the observation of biomodal timing distribution of Aneu-OGs, compared with the unimodal background genes, supports the interpretation that a minority of Aneu-OGs function as drivers, while most of them that occurs distinctly later likely act as enablers of aneuploidy tolerance, with CCT5 being a notable example discussed below. Notably, no Aneu-OGs fell into the late timing indices (Fig. 4C). This depletion indicates that once tumor cells have acquired sufficient Aneu-OGs to proliferate and survive in the aneuploid state, additional gains confer limited incremental benefit, and genes with terminal timing indices are therefore not typically selected after aneuploidy tolerance has been established.

This interpretation is supported by the observation that gains of 1q and 8q, which our analysis classifies as relatively early events, are consistently enriched across cancer types. The early timing of 1q gain is particularly well documented in breast cancer (**Fig. S2C**) and other adenocarcinomas, including liver and gastric cancers. Single cell DNA sequencing of healthy breast tissue has shown that 1q gain is already present in rare, clonally expanded epithelial cells of healthy women, before any cancer develops, and accumulates with age [41]. Similar precancerous 1q gained populations have been detected in luminal breast epithelial cells from both BRCA1/BRCA2 mutation carriers and wild-type individuals, often arising before LOH at the BRCA loci [42]. Functional screens further establish 1q gain as a *bona fide* driver event in breast tumorigenesis: forward genetic evolution experiments in untransformed mammary epithelial cells show that 1q gain confers a proliferative advantage through Notch signaling potentiation by 1q resident γ-secretase genes, recapitulating the tissue specific selection pattern observed in patient tumors [19]. Independent timing analyses based on whole genome sequencing in the Pan-Cancer Analysis of Whole Genomes (PCAWG) cohort similarly identified specific copy number gains as early oncogenic events, including trisomy 7 in glioblastoma and isochromosome 17q in medulloblastoma, while broader genomic instability emerged as a feature of later tumor evolution [43], consistent with our finding that Aneu-OGs are selected early in tumor evolution.

The pathways enriched among early Aneu-OGs further support their assignment as drivers. CC regulation and mitotic regulation, including the G1/S transition, G1 phase, and mitotic anaphase, were consistently among the earliest enriched pathways across cancer types. This is mechanistically coherent, since dysregulation of CC progression and mitotic fidelity is a direct cause of chromosome missegregation and, therefore, of aneuploidy itself. The early selection of gains affecting these pathways is consistent with a model in which the initial establishment of aneuploidy is driven by alterations that increase missegregation rates, rather than by alterations that merely permit the survival of already aneuploid cells. The late timing of protein folding pathways, including the CCT/TRiC chaperonin complex that contains CCT5, fits their complementary role of enablers that buffer the proteotoxic stress of the aneuploid state.

The majority of Aneu-OGs occurring later relatively to the all set of Aneu-OGs likely represents genuine enablers of aneuploidy tolerance. Although aneuploidy is typically viewed as a catastrophic cellular state that triggers apoptosis in normal cells, the predominance of late-occurring Aneu-OGs suggests that cancer progression depends less on the initial generation of CIN and more on acquiring the capacity to survive it. This is consistent with a recent study of karyotypic adaptation following spindle assembly checkpoint inhibition [44], in which the recurrent aneuploidies that emerged across cell lines, including gains of chromosome arms 1q, 3q, and chromosomes 5, 8, and 20, were highly concordant with our pan-cancer Aneu-OG distribution. The repeated selection of these specific aneuploidies during experimental adaptation to chronic CIN supports the conclusion that COATS captures selected and biologically meaningful events. A recent study further demonstrated that cells can develop an oncogene-like addiction to specific aneuploidies, with these chromosomal gains becoming selectively indispensable for continued proliferation [45]. The abundance of aneuploidy enablers relative to early drivers also carries evolutionary implications. These tolerance mechanisms broaden the range of karyotypic states that remain viable, increasing both the genomic diversity available within tumor cell populations and the probability of arising rare variants conferring resistance to therapy or enable survival during metastatic spread [46]. Cancer therefore behaves not as a static collection of mutations, but as a dynamically evolving population in which CIN generates diversity and selection determines which variants persist [47].

### Aneu-OGs are preferentially selected early during tumor evolution compared with cell cycle or immune evasion

Compared with genome wide timing events, Aneu-OGs showed a highly recurrent pattern of enrichment in early events and depletion in late events (21 cancer types or groups, Table S3), whereas this pattern was substantially less frequent for CC (13 cancer types or groups, Table S3) and immune related oncogenes (9 cancer types or groups, Table S3). This contrast suggests that the early timing signal is not a generic feature of proliferation or immune evasion in cancer, but is more specific to oncogenes linked to aneuploidy. We note that this gene level contrast is distinct from the pathway level timing described above: although CC and mitotic pathways are among the earliest enriched pathways within the Aneu-OG set, the broader collection of CC oncogenes defined genome wide does not show the same recurrent early enrichment, indicating that early selection is specific to CC and mitotic genes that are also linked to aneuploidy, e.g. CCNE2 (**Fig. 4C**), rather than to CC genes in general. The depletion of Aneu-OGs in late events indicates that once aneuploidy tolerant cells have emerged, additional gains of these genes may confer a smaller incremental advantage.

### CCT5, a COATS-identified Aneu-OG, reveals a vulnerability of aneuploid cells

The experimental validation of CCT5 illustrates both the biological logic of aneuploidy tolerance and its potential therapeutic relevance. CCT5 encodes a subunit of the chaperonin-containing TCP-1 complex, whose role in folding cytoskeletal proteins and numerous other essential substrates makes it particularly critical when the proteotoxic burden is elevated. Aneuploid cells experience chronic protein folding stress arising from stoichiometric imbalances across protein complexes, and aneuploid yeast strains show consistent upregulation of CCT/TRiC complex subunits as a direct response to this burden [39]. The late timing of 5p gain, on which CCT5 resides, classifies it as a tolerance enabler rather than an early driver of CIN, consistent with a model in which cells that have already acquired an aneuploid karyotype subsequently select for amplification of protein quality control machinery. Our multicolor competition assay confirms that CCT5 depletion selectively impairs the competitive fitness of aneuploid HCEC clones while leaving the near-diploid clones unaffected. CCT complex subunits have previously been shown to be overexpressed across multiple cancer types and associated with poor prognosis in patients [48–50], and CCT2 depletion has been shown to prevent tumor growth in a syngeneic breast cancer model [51], establishing a functional precedent for targeting individual CCT subunits as a therapeutic strategy for cancer. Our findings add a specific mechanistic rationale for CCT5 essentiality in the aneuploid context, demonstrating that this dependency is not a general cancer cell property but one that is selectively heightened by karyotypic imbalance. The co-alteration of multiple CCT complex subunits across cancer types (Fig. 5E) further suggests that the complex as a whole may warrant investigation as a therapeutic target in chromosomally unstable cancers. More broadly, resensitizing tumor cells to the stresses imposed by their own genomic instability, rather than attempting to block instability at its source, represents a conceptually distinct and potentially complementary therapeutic strategy for cancers characterized by high CIN and aneuploidy [25, 38]. Because aneuploid cancer cells already operate near the limits of proteotoxic and transcriptional tolerance, a further reduction of chaperonin activity is selectively lethal to these cells while normal diploid cells remain comparatively insensitive, defining a context-specific vulnerability that is in principle exploitable across the broad spectrum of cancers in which late-timing copy number gains of protein quality control genes are selected.

## Materials and Methods

### Data preprocessing

Datasets of all 33 types of cancer in TCGA, including Acute Myeloid Leukemia (LAML), Adrenocortical carcinoma (ACC), Bladder Urothelial Carcinoma (BLCA), Brain Lower Grade Glioma (LGG), Breast invasive carcinoma (BRCA), Cervical squamous cell carcinoma and endocervical adenocarcinoma (CESC), Cholangiocarcinoma (CHOL), Colon adenocarcinoma (COAD), Esophageal carcinoma (ESCA), Glioblastoma multiforme (GBM), Head and Neck squamous cell carcinoma (HNSC), Kidney Chromophobe (KICH), Kidney renal clear cell carcinoma (KIRC), Kidney renal papillary cell carcinoma (KIRP), Liver hepatocellular carcinoma (LIHC), Lung adenocarcinoma (LUAD), Lung squamous cell carcinoma (LUSC), Lymphoid Neoplasm Diffuse Large B-cell Lymphoma (DLBC), Mesothelioma (MESO), Ovarian serous cystadenocarcinoma (OV), Pancreatic adenocarcinoma (PAAD), Pheochromocytoma and Paraganglioma (PCPG), Prostate adenocarcinoma (PRAD), Rectum adenocarcinoma (READ), Sarcoma (SARC), Skin Cutaneous Melanoma (SKCM), Stomach adenocarcinoma (STAD), Testicular Germ Cell Tumors (TGCT), Thymoma (THYM), Thyroid carcinoma (THCA), Uterine Carcinosarcoma (UCS), Uterine Corpus Endometrial Carcinoma (UCEC), Uveal Melanoma (UVM), were used in this paper.

We downloaded harmonized TCGA gene expression data processed through HTSeq-FPKM (High-Throughput Seq-Fragments Per Kilobase of transcript per Million mapped reads) workflow, copy number segment (CNS) data and allele-specific copy number segment data from Genomic Data Commons using the *TCGAbiolinks* package [52]. We downloaded the aneuploidy scores from the Pan-Cancer Atlas study [3]. The tumor purity was obtained from ABSOLUTE [15]. The publicly available MC3 (the Multi-Center Mutation Calling in Multiple Cancers) data [22] were downloaded directly from Genomic Data Commons. The MC3 data were generated under GRCH37, and all mutations were lifted to GRCH38 using R package *rtracklayer* [53] to match with the reference genome of SCNA data.

The expression data are log2 transformed. 56830 genes are left and used in this paper after genes having 0 value across all samples from any type of cancer were excluded from the whole datasets. Gene-level SCNA values were inferred from the corresponding CNS data. If a gene does not fall into any segment in CNS data, then its SCNA value was inferred by the mean CNS value of two adjacent segments.

### Association Sign

The cancer-specific association sign between two variables is given by the consistent sign observed in both Pearson and Spearman correlation coefficients. If these two coefficients do not align, the cancer-specific association sign is considered ambiguous. The multi-cancer association sign is defined by the consistent sign of the weighted median of Pearson and Spearman correlations computed across these cancer types where the weights are given by the corresponding numbers of samples. In instances where these correlations do not present a consistent sign, the multi-cancer association sign is considered ambiguous.

### Association function and statistical significance

We used R package *conMItion* [16] for estimating the association and the corresponding statistical significance. Normalized Mutual Information (MI) is used to measure the cancer-specific association between gene expression and SCNA (expression-SCNA association) for each gene. Normalized conditional Mutual Information (CMI) is used to measure the cancer-specific expression-aneuploidy association and SCNA-aneuploidy association given the condition of tumor purity. The multi-cancer expression-SCNA, expression-aneuploidy, or SCNA-aneuploidy association for a gene is given by the weighted median of the corresponding cancer-specific associations, where the weights are given by the sample numbers of the corresponding cancer types.

For each cancer type, a null distribution size of 10,000,000 is generated by the *conMItion* package. To generate the multi-cancer distribution, we calculate the weighted median of the sorted cancer-specific distributions across different cancer types. The P-value for cancer-specific (or multi-cancer) associations is determined by its rank within the respective cancer-specific (or multi-cancer) distributions. False discovery rates are used for multiple hypothesis testing correction. Associations with a Q value below 0.2 are considered statistically significant.

### Workflow of COATS

MI and CMI are used to estimate association in this paper. Then permutation-based tests are used to evaluate the statistical significance of the three types of association. The proposed method selects cancer-specific/pan-cancer amplification-dependent (deletion-dependent) OGs (TSGs) based on the following 3 criteria: (a) genes whose expression levels are significantly and positively (negatively) associated with the AS, (b) genes whose SCNA levels are significantly and positively (negatively) associated with the AS, and (c) genes whose expression levels are significantly associated with their SCNA levels.

### Statistical significance of shared OGs/TSGs

We collected the number of OGs and TSGs for each cancer type or cancer group. The ratio of these counts to the total number of genes gives the probability of a random gene being an OG or TSG in a specific cancer type or group. This scenario can be modeled using a Poisson binomial distribution, treating each cancer type as an independent Bernoulli trial. We can calculate the probability of a gene being an OG or TSG in more than a certain number of cancer types or groups. Subsequently, the P value of observing a specified number of such genes across different cancer types can be determined based on this binomial distribution. The P-value for observing a specified range of genes across various cancer types is calculated by summing the P-values for each gene count. In this paper, the range of genes are given by ±5 of the observed OGs and TSGs.

### Timing index of identified OGs

We employed a comparative strategy between SCNA and mutation [23, 24] to infer whether an OG is an early or late event. For each cancer sample, we assigned a timing index to every copy number segment (CNS). For a given segment, we identified all loci with mutations from the MC3 data. We then determined the copy number containing these mutations for each locus within the segment based on mutation and reference allele counts in MC3 data. If this number is less than the copy number of the major allele of the segment, it suggests that the SCNA of this segment occurred earlier than the mutation at the given locus. The timing index is calculated as the percentage of loci within the segment where the SCNA is identified as occurring earlier than the mutations. Only amplifications are included in the timing index calculation and deletions are removed from the analysis. The gene-level timing index can be inferred in the similar way to deriving gene-level SCNA values from CNS data. The timing index for a gene in a specific type of cancer is given by the mean timing index across all samples. The multi-cancer timing index is given by the weighted median of the timing indices from the corresponding cancer types.

### Timing index of REACTOME pathways

We also assigned a timing index to each identified enriched REACTOME pathway. Th index of a pathway is given by the average timing index of the identified OGs that overlap with this pathway.

### Simulation of modeling chromosome-arm gains as CIN enablers and drivers

We examined two different scenarios for how the copy number aberrations affecting one chromosome arm may impact cancer evolution, by modifying CINner [21], a simulator for CIN. In both scenarios, every time a cell divides, a random chromosome arm is missegregated with probability *p_misseg_*. We define each cell’s CIN score to be the total number of arm missegregations it has acquired. Given a cell with copy number *c_k_* of arm *k*, its fitness equals 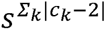. A cell is nonviable and dies if it suffers from nullisomy (i.e., *c_k_* =0 for some arm *k*) or its CIN score exceeds limit *L_CIN_*. Otherwise, the cell expands at a rate corresponding to its fitness.

The first scenario (namely “CIN enabler”) models the case where CIN becomes more tolerated with gains of a chromosome arm, e.g. 5*p*. Here, the missegregation rate *p_misseg_* = 2.5 • 10^-5^ is constant, but the viability limit *L_CIN_* = 5(1 + 3 • *max*(0, *c*_5_*_p_* - 2)) grows as the CN of arm 5*p* increases.

The second scenario (namely “CIN driver”) models the case where CIN rates depend on arm 5*p*. In this scenario, the viability limit *L_CIN_* = 45 is the same for all cells, but each cell’s missegregation rate *p_misseg_* = 1.5 • 10^-6^(1 + 20 • *max*(0, *c*_5_*_p_* - 2)) increases with gains of arm 5*p*.

The parameters for each scenario are selected such that the CIN scores are similar for different *c*_5_*_p_*. In both cases, we choose selection strength *s*= 3. 10,000 simulations are created for each scenario. For each simulation, we define the “normalized true time” that a 5*p* event occurs to be 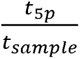, where *t*_5*p*_ is the time that the event occurs and *t_sample_* is the time where the CN profile is sampled from the simulation. We also define the event’s “normalized CN time” to be 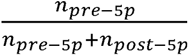, where *n_pre−5p_* and *n_post−5p_* are the number of chromosome-arm missegregations occurring before and after the 5*p* event, respectively.

### Western Blot Analysis

For Western blot analysis, cells were harvested by trypsinization, washed with cold PBS, and lysed in RIPA lysis buffer (Abcam, ab156034) supplemented with protease and phosphatase inhibitor cocktail (Thermo Scientific, 78441). Lysates were agitated for 30 minutes at 4°C, followed by centrifugation at 4°C. Supernatants were collected and mixed with Pierce LDS buffer (Thermo Scientific, 84788) and Bond-Breaker TCEP solution (Thermo Scientific, 77720) at concentrations recommended by the manufacturers. Lysates were then boiled for 10 minutes at 80°C.

After returning to room temperature, lysates were loaded into wells of NuPAGE 4-12% Bis-Tris mini gels (Thermo Scientific, NP0323BOX) for electrophoresis, followed by transfer to PVDF membranes (Bio-Rad, 1704274) using a semi-dry transfer system (Trans-Blot Turbo transfer system, Bio-Rad, 1704150). Membranes were blocked with 5% non-fat milk in Tris-buffered saline (TBS) with 0.1% Tween-20 (TBST) for 1 hour at room temperature, followed by overnight incubation with primary antibodies diluted in 2.5% BSA in TBS at 4°C.

The following primary antibodies were used: CCT5 (Abcam) at a dilution of 1:1000, and GAPDH (Santa Cruz, sc-47724) at a dilution of 1:10,000. Membranes were washed three times with TBST and incubated with IRDye 680RD goat anti-mouse IgG secondary antibody (926-68070) at a dilution of 1:20,000 or IRDye 800CW goat anti-rabbit IgG secondary antibody (926-32211) at a dilution of 1:20,000 for 2 hours at room temperature. After three washes with TBST, signals were detected using the Li-Cor Odyssey DLx imaging system. Quantification of Western blot images was performed using ImageJ by calculating the signal derived from the indicated bands and subtracting the relative background, with normalization performed relative to GAPDH as the internal control protein.

### shRNA-Mediated Knockdown

For CCT5 knockdown, five different shRNAs targeting human CCT5 were obtained with the following target sequences:

- CCT5 shRNA-1 (sh1): CCGGCCGGGTTATTGTGTGTAACTACTCGAGTAGTTACACACAATAACCCGGTTTTTG
- CCT5 shRNA-2 (sh2): CCGGTCTTCAATAATGGCAGCAACTTCTCGAGAAGTTGCTGCCATTATTGAAGATTTTTTG
- CCT5 shRNA-3 (sh3): CCGGCAACTTCTGATTGATTACACACTCGAGTGTGTAATCAATCAGAAGTTGTTTTTG
- CCT5 shRNA-4 (sh4): CCGGGACGAAACATCCGGATTAATTCTCGAGAATTAATCCGGATGTTTCGTCTTTTTG
- CCT5 shRNA-5 (sh5): CCGGCTGCAAACATTGATAGGTCATCTCGAGATGACCTATCAATGTTTGCAGTTTTTG

The MISSION® pLKO.1-puro non-mammalian shRNA plasmid targeting no known mammalian genes was used as the control shRNA (Sigma-Aldrich, SHC002).

Lentiviral particles were produced by co-transfecting HEK293T cells with shRNA plasmids along with lentiviral packaging and envelope plasmids. Target cells were transduced with the lentivirus and selected with puromycin to generate stable knockdown cell lines.

### Multicolor Competition Assay

To quantitatively assess the differential requirement for CCT5 in near-diploid versus aneuploid cells, we performed a multicolor competition assay. Near-diploid and aneuploid HCEC clones were transduced with either control shRNA or CCT5 shRNA-3. For each comparison, one population was fluorescently labeled with GFP while the other population remained unlabeled. Labeled and unlabeled cells were mixed at an initial 1:1 ratio based on cell counts and co-cultured in standard culture medium for 7-10 days to allow multiple rounds of cell division.

After the co-culture period, cells were collected by trypsinization and the relative proportions of labeled (GFP-positive) and unlabeled populations were quantified by flow cytometry using a SONY SH800 cell sorter. To control for potential effects of fluorescent protein expression on cell fitness, each experiment was performed in triplicate with fluorescent labels switched between the control shRNA and CCT5 shRNA populations across replicates.

Relative viability was calculated as the ratio of CCT5 shRNA-expressing cells to control shRNA-expressing cells at the end of the competition period, normalized to the initial 1:1 mixing ratio. A relative viability value less than 1 indicates that CCT5 knockdown impairs competitive fitness. Statistical significance was assessed using the Wilcoxon rank-sum test, with P < 0.01 considered significant. Data are presented as mean ± standard deviation from three independent experiments.

## Code Availability

All the R codes used in COATS can be downloaded at GitHub repository https://github.com/GJYWang/COATS.

## Data Availability

All the data used in COATS are publicly available. Access of these data has been described in Materials the Methods section. Processed data used to run COATS can be accessed through https://owww.molgen.mpg.de/~COATS/COATS.tar.gz.

## Acknowledgments

We thank the valuable discussion with Dmitri A. Petrov. We thank all members of the Davoli laboratory for helpful discussions. This research was supported by NIH R37CA248631, R01HG012590, R01DK135089 (T.D.), R01CA310281 (K.N.D.), and the Pershing Square Sohn Cancer Research Award (T.D.).

## Supplementary Figures

**Fig. S1.**
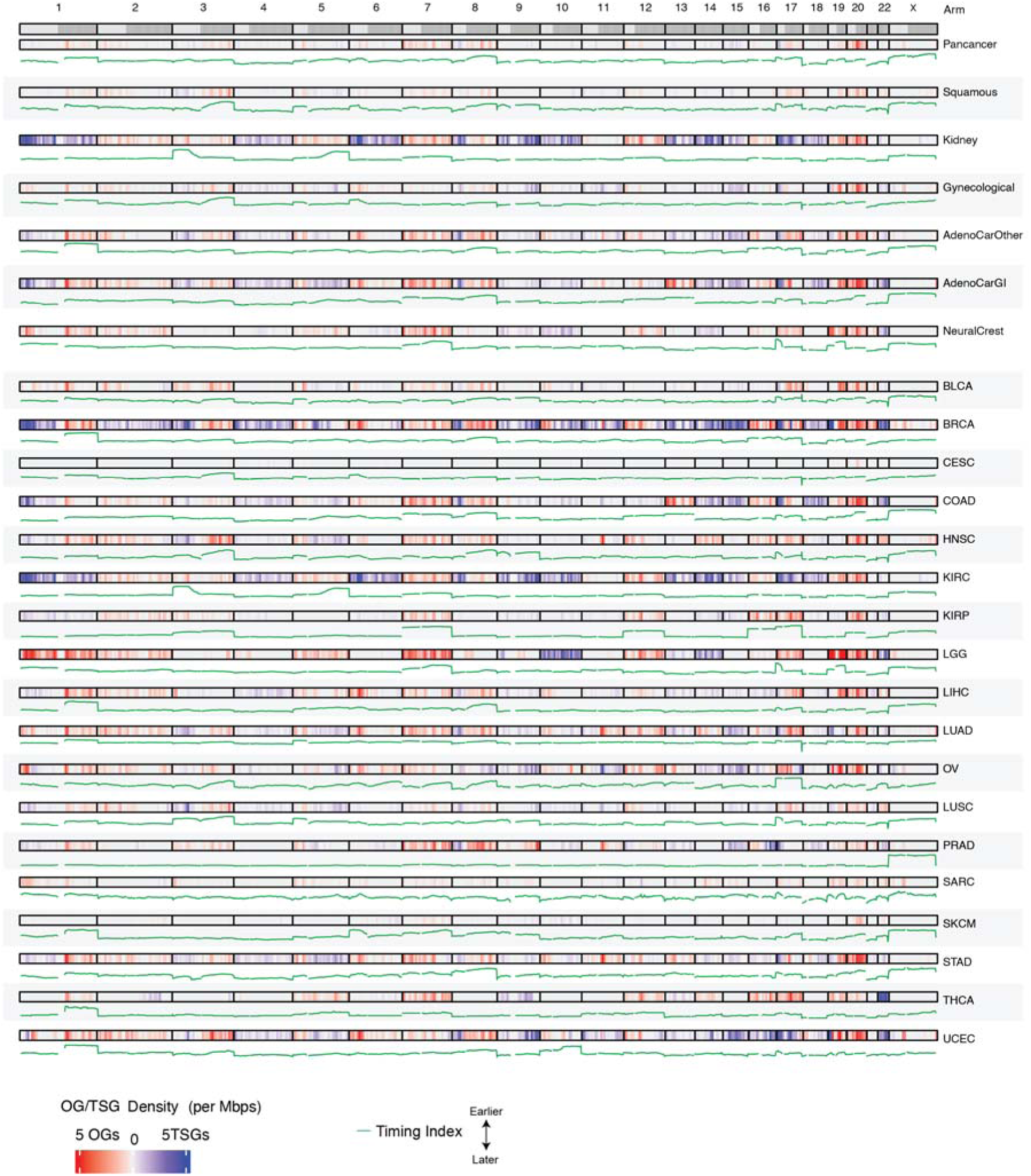
Genome-wide distribution of Aneu-OGs, Aneu-TSGs, and timing index across cancer groups and cancer types. Each row represents a cancer group or cancer type. Aneu-OG and Aneu-TSG densities were calculated in 1 Mb genomic windows and plotted across chromosome arms (1–22 and X). Red indicates Aneu-OG density and blue indicates Aneu-TSG density; white indicates no Aneu-OGs or Aneu-TSGs. The timing index (green line) is shown below each density track to illustrate its genome-wide variation relative to Aneu-OG and Aneu-TSG densities.

**Fig. S2.**
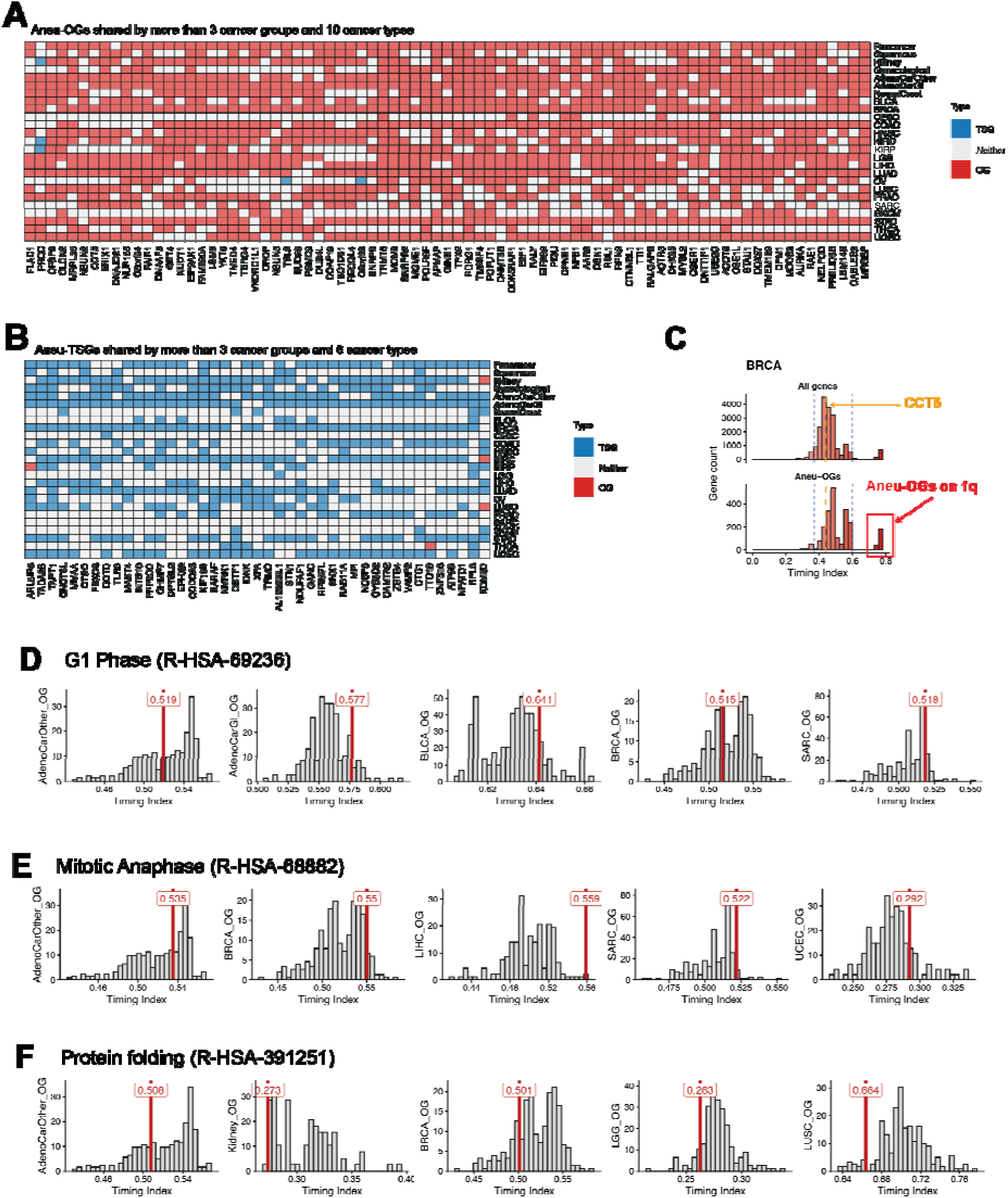
Aneu-OGs and Aneu-TSGs shared across cancer types and cancer groups, and timing of representative pathways. **(A)** Aneu-OGs shared across cancer types and cancer groups. Each row of the heatmap represents a cancer type or group, and each column corresponds to a gene classified as an Aneu-OG (red), Aneu-TSG (blue), or neither (white). The heatmap shows Aneu-OGs shared by more than 3 cancer groups and 10 cancer types (18 major cancer types included). Aneu-OGs are frequently shared across cancer types. **(B)** Aneu-TSGs shared across cancer types and cancer groups. Each row represents a cancer type or group, and each column corresponds to a gene classified as an Aneu-TSG (blue), Aneu-OG (red), or neither (white). The heatmap shows Aneu-TSGs shared by more than 3 cancer groups and 5 cancer types (18 major cancer types included). Aneu-TSGs are more tissue specific than Aneu-OGs. **(C)** Timing index distributions for all genes compared with Aneu-OGs across in BRCA. The upper histogram shows the unique timing indices of all genes and the lower histogram shows the timing indices of Aneu-OGs. The timing indices were classified as early, late, or terminal events using the 5th and 95th percentile thresholds of the timing indices of all the genes, indicated by the blue dashed lines. The timing index for CCT5 (orange) and Aneu-OGs on 1q (red) are marked. **(D)** Timing index (red vertical line) of the Reactome pathway G1 Phase (R-HSA-69236) plotted against the histogram of timing indices for all enriched Aneu-OG pathways in selected cancer types or cancer groups. **(E)** Timing index (red vertical line) of the Reactome pathway Mitotic Anaphase (R-HSA-68882) plotted against the histogram of timing indices for all enriched Aneu-OG pathways in selected cancer types or cancer groups. **(F)** Timing index (red vertical line) of the Reactome pathway Protein folding (R-HSA-391251) plotted against the histogram of timing indices for all enriched Aneu-OG pathways in selected cancer types or cancer groups.

**Fig. S3.**
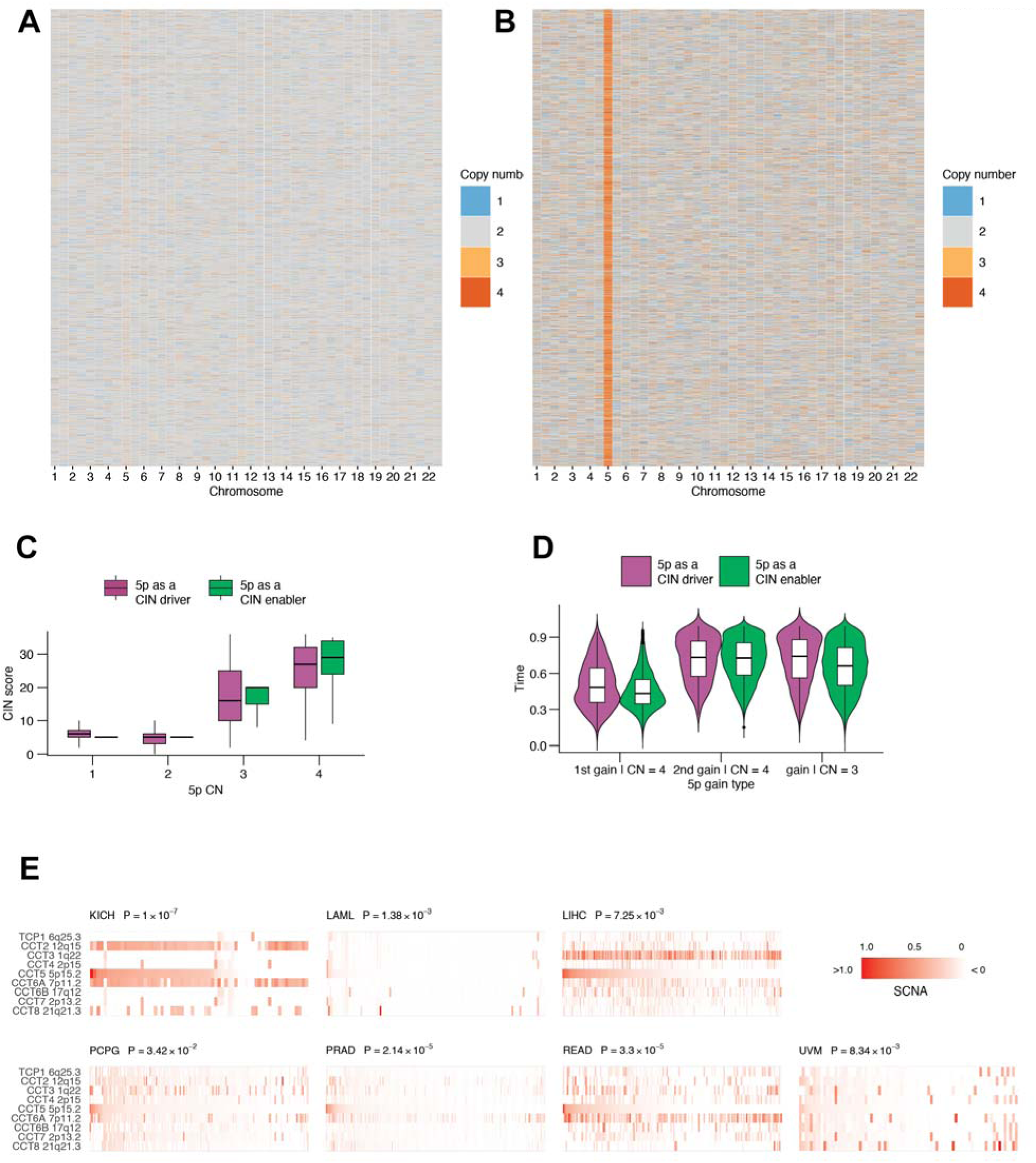
CINner simulations of 5p as CIN driver versus enabler, and supporting analyses of the CCT gene family. **(A)** Copy number profiles of samples simulated under the assumption that 5p SCNAs act as aneuploidy drivers (schematic in Fig. 4A). **(B)** Copy number profiles of samples simulated under the assumption that 5p SCNAs act as aneuploidy enablers (schematic in Fig. 4A). In A and B, each row corresponds to a synthetic sample, each column to an arm, with color indicating copy number. **(C)** Distribution of CIN scores as a function of arm 5p CN, compared between the two scenarios. **(D)** Distribution of real-times when SCNAs on arm 5p occur, compared between the two scenarios. SCNAs are grouped into three categories: exclusive 5p gains resulting in observed CN = 3, and first or second 5p gains resulting in observed CN = 4. **(E)** The heatmaps of the SCNAs for the CCT family genes across KICH, LAML, LIHC, PCPG, PRAD, READ, and UVM. Amplification is visualized using a color scale indicating amplification (red), with color intensity representing the degree of amplification. SCNA values below 0 were colored in white. Gene names and their respective cytogenetic locations are labeled on the left side of the heatmap.

**Fig. S4.**
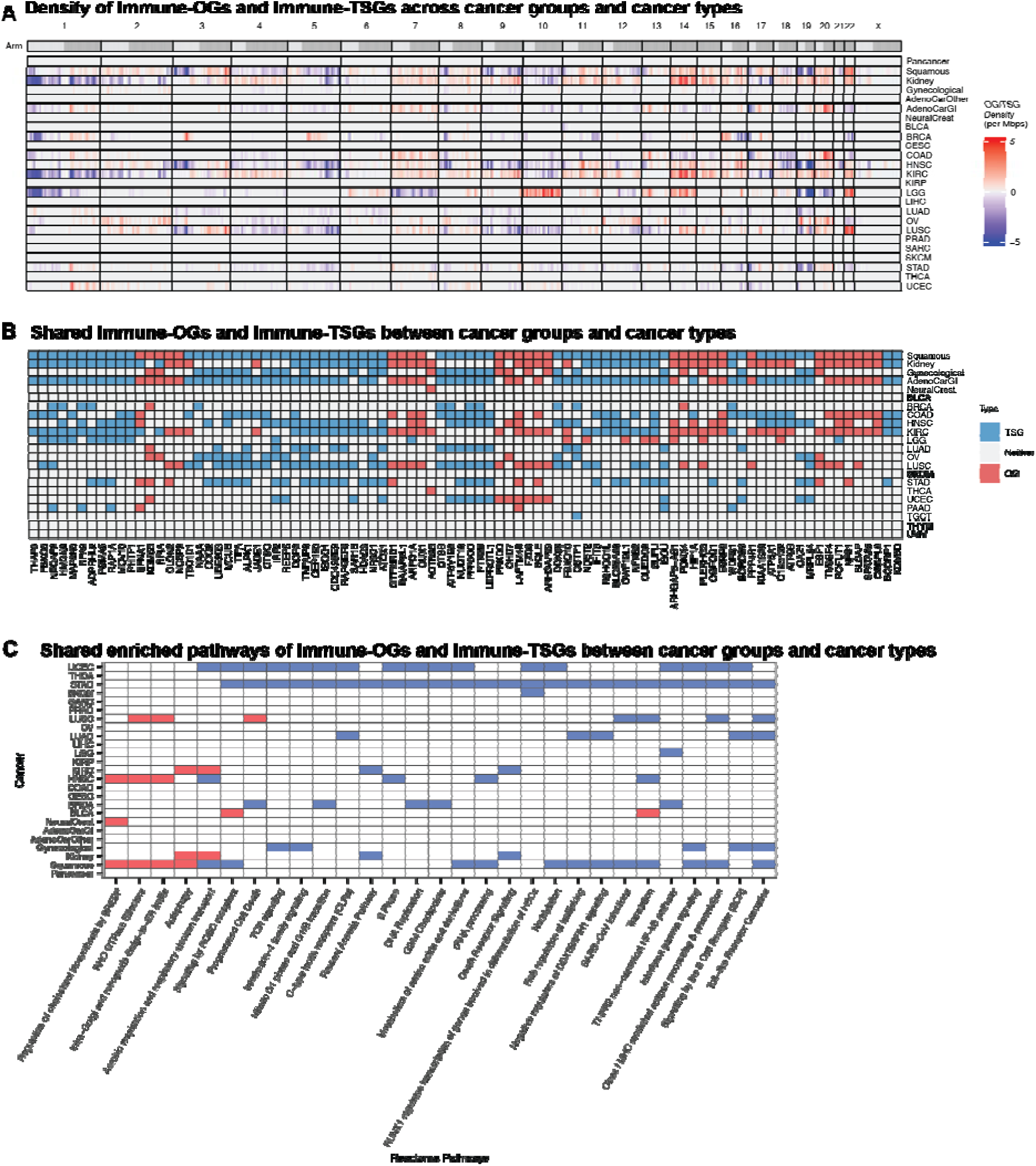
Immune-OGs and Immune-TSGs across cancer types and cancer groups. **(A)** Genome-wide distribution of Immune-OGs and Immune-TSGs across cancer groups and cancer types. Each row represents a cancer type or cancer group, and each column corresponds to a chromosome arm (1–22 and X). The color gradient indicates the density of Immune-OGs and Immune-TSGs per Mb, with red indicating Immune-OG density and blue indicating Immune-TSG density; white indicates no Immune-OGs or Immune-TSGs. **(B)** Immune-OGs and Immune-TSGs shared across cancer types and cancer groups. Each row represents a cancer type or cancer group, and each column corresponds to a gene classified as an Immune-OG (red), Immune-TSG (blue), or neither (white). The heatmap shows Immune-OGs and Immune-TSGs shared by more than 2 cancer groups (18 major cancer types included). **(C)** Heatmap of enriched pathways shared across cancer types and cancer groups. Each row represents a cancer type or cancer group, and each column represents a Reactome pathway. Red cells indicate pathways enriched among Immune-OGs, blue cells indicate pathways enriched among Immune-TSGs, and white cells indicate pathways with no significant enrichment.

**Fig. S5.**
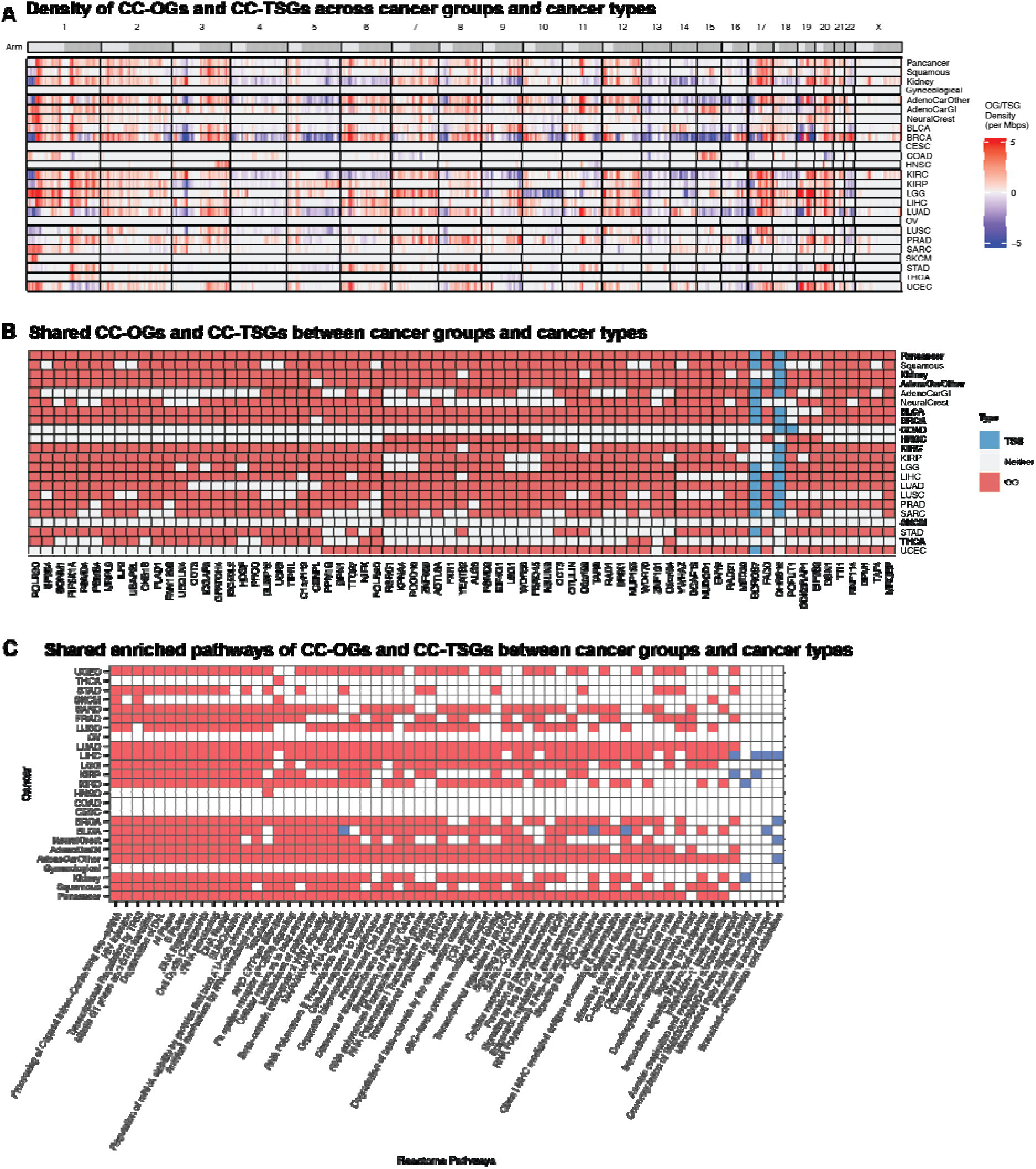
CC-OGs and CC-TSGs across cancer types and cancer groups. **(A)** Genome-wide distribution of CC-OGs and CC-TSGs across cancer groups and cancer types. Each row represents a cancer type or cancer group, and each column corresponds to a chromosome arm (1–22 and X). The color gradient indicates the density of CC-OGs and CC-TSGs per Mb, with red indicating CC-OG density and blue indicating CC-TSG density; white indicates no CC-OGs or CC-TSGs. **(B)** CC-OGs and CC-TSGs shared across cancer types and cancer groups. Each row represents a cancer type or cancer group, and each column corresponds to a gene classified as a CC-OG (red), CC-TSG (blue), or neither (white). The heatmap shows CC-OGs and CC-TSGs shared by more than 3 cancer groups and 10 cancer types (18 major cancer types included). **(C)** Heatmap of enriched pathways shared across cancer types and cancer groups. Each row represents a cancer type or cancer group, and each column represents a Reactome pathway. Red cells indicate pathways enriched among CC-OGs, blue cells indicate pathways enriched among CC-TSGs, and white cells indicate pathways with no significant enrichment.

